# Xela DS2 and Xela VS2: two novel skin epithelial-like cell lines from adult African clawed frog (*Xenopus laevis*) and their response to an extracellular viral dsRNA analogue

**DOI:** 10.1101/2020.05.08.084723

**Authors:** Maxwell P. Bui-Marinos, Joseph F. A. Varga, Nguyen T.K. Vo, Niels C. Bols, Barbara A. Katzenback

## Abstract

The skin epithelial layer acts as an important immunological barrier against pathogens and is capable of recognizing and responding to pathogen associated molecular patterns (PAMPs) in human and mouse models. Although presumed, it is unknown whether amphibian skin epithelial cells exhibit the ability to respond to PAMPs such as viral double-stranded RNA (dsRNA). To address this, two cell lines from the dorsal skin (Xela DS2) and ventral skin (Xela VS2) of the African clawed frog (*Xenopus laevis*) were established. Xela DS2 and Xela VS2 cells have an epithelial-like morphology, express genes associated with epithelial cells, and lack senescence-associated beta-galactosidase activity. Cells grow optimally in 70% Leibovitz’s L-15 medium supplemented with 15% fetal bovine serum at 26°C. Upon treatment with poly(I:C), a synthetic viral dsRNA analogue and known type I interferon inducer, Xela DS2 and Xela VS2 exhibit marked upregulation of key pro-inflammatory and antiviral transcripts suggesting frog epithelial cells participate in the recognition of extracellular viral dsRNA and production of local inflammatory signals; similar to human and mouse models. Currently, these are the only known *Xenopus laevis* skin epithelial-like cell lines and will be important for future research in amphibian epithelial cell biology, initial host-pathogen interactions, and rapid screening of the effects of environmental stressors, including contaminants, on frog skin epithelial cells.

## 1. Introduction

Amphibian skin is critical for homeostatic physiological functions, albeit species-dependent, such as osmoregulation, respiration, and ion transport (Haslam et al., 2014; Lillywhite, 2006), and immune defence against pathogens (Stuart et al., 2004). Amphibian skin innate immune functions comprise of a physical barrier formed through junctions between epithelial cells, a chemical barrier through the synthesis and secretion of antimicrobial peptides from specialized glands into the skin mucous layer, and a cellular barrier comprised of many cell types that participate in the recognition and response to pathogens (Varga et al., 2019). While frog antimicrobial peptides and their direct antimicrobial activities to pathogens are of human and amphibian importance (Rollins-Smith, 2009), relatively little investigation has focused on amphibian skin epithelial cell innate immune function. Amphibian skin tissue is complex in cellular composition, including epithelial cells in the outer epidermal layer, mainly fibroblasts in the underlying dermal layer, and interdigitating resident immune cells throughout the skin such as Langerhans cells and mast cells (Haslam et al., 2014). As frog skin epithelial cells are in direct and constant contact with the external environment, understanding the ability of these cells to respond to pathogens and initiate innate immune responses is crucial to understanding the complex host-pathogen interactions occurring at the skin interface.

Recent evidence from murine and human models suggest that epithelial cells are more than just bystanders, playing an active role in responding to pathogens and downstream communication to neighbouring cells, including immune cells (Hamel et al., 2015; Pasparakis et al., 2014). Like other cell types, epithelial cells express a number of pattern recognition receptors (PRRs) that sense pathogen-associated molecular patterns (PAMPs), leading to the activation of key transcription factors (interferon regulatory factor 3, IRF3/3; nuclear factor kappa light chain enhancer of activated B cells, NF-kB, activator protein-1, AP-1). The recognition of select PAMPs activates transcription factors involved in the regulation of antiviral and proinflammatory gene expression; ultimately controlling cytokines and chemokines, at the transcriptional level, during the immune response. (Artis, 2008; Ghosh et al., 1998; Tosi, 2005). During viral replicative cycles, intracellular viral double-stranded RNA (dsRNA) is produced (Weber et al., 2006) and is detected as a viral PAMP by the retinoic acid-inducible gene I (RIG-I) receptor and melanoma differentiation-associated protein 5 (MDA5) in the cytosol. Extracellular viral dsRNA, likely released as a result of host cell lysis, is trafficked to endosomes by class A scavenger receptors [SR-As; (DeWitte-Orr et al., 2010)], where it is recognized by endosomal toll-like receptor 3 [TLR3; (DeWitte-Orr et al., 2010; DeWitte-Orr et al., 2007)]. Recognition of extracellular viral dsRNA by TLR3 results in the rapid activation of downstream signaling cascades, culminating with the translocation of homodimeric IRF3 and NF-κB that together mediate the expressions of type I interferons (Type I IFNs), proinflammatory cytokines and chemokines (Artis, 2008; DeWitte-Orr et al., 2010; Ghosh et al., 1998; Tosi, 2005). Secreted type I IFNs can then act in an autocrine or paracrine fashion through their cognate IFN receptors (IFNARs) on neighbouring cells to initiate antiviral programs through signal transducer and activator of transcription (STAT) and interferon regulatory factor (IRF) proteins. Binding of IFNAR1/2 leads to the activation of the transcription factor complex ISGF3, comprised of STAT1/STAT2 and IRF9, and binds interferon stimulated response elements (ISREs) to regulate the expression of interferon stimulated genes (ISGs). Many of these ISGs encode for the proteins necessary for mediating antiviral programs in the host (Fink and Grandvaux, 2013; Schoggins and Rice, 2011) and includes IRF7, a transcription factor that can heterodimerize with IRF3 to further bolster the induction of type I IFNs in a positive feedback loop (Ysebrant de Lendonck et al., 2014). Some of the most well characterized dsRNA-induced antiviral genes include myxovirus resistance protein family (MX) and double-stranded RNA dependent protein kinase R (PKR) (Schoggins and Rice, 2011).

Despite the documented roles of skin epithelial cells in pathogen sensing and initiation of proinflammatory and antiviral responses in mouse and human models (Hamel et al., 2015; Nestle et al., 2009; Pasparakis et al., 2014), very little is known regarding whether frog skin epithelial cells function in a similar fashion. Among frog species studied, the African clawed frog (*Xenopus laevis*) has been established as a model system to investigate host-pathogen interactions and amphibian immunity (Gantress et al., 2003). However, the contribution of isolated skin epithelial cells to amphibian antiviral immunity remains limited. It is critical to dissect the potential individual role(s) of frog skin epithelial cells in regard to responding to viral PAMPs to improve our understanding of the skin epithelial cell antiviral mechanisms. Since dsRNA is produced by the majority of viruses at some point during their replicative life cycle (Weber et al., 2006), the addition of poly(I:C), a viral dsRNA analogue and a known inducer of Type I IFN (Poynter and DeWitte-Orr, 2015), may mimic extracellular viral dsRNA that is released from lysed infected cells. The objectives of this study were to first generate and characterize novel *X. laevis* skin epithelial cell lines and then use these newly generated frog skin cell lines as a model system to elucidate the role of frog skin epithelial cells in responding to dsRNA and initiating antiviral gene expression programs. We hypothesized that these frog skin cell lines would upregulate proinflammatory and antiviral cytokine transcript levels in response to extracellular poly(I:C). Herein, we report on the generation and establishment of two skin epithelial-like cell lines from *X. laevis* dorsal and ventral skin, named Xela DS2 and Xela VS2, respectively, their optimal growth conditions, morphological and molecular characterization, and their capacity to initiate antiviral gene expression pathways in response to poly(I:C).

## 2. Methods

### 2.1. Frogs

Dorsal and ventral skin tissues from adult female African clawed frogs (*Xenopus laevis*) were kindly donated by Dr. Mungo Marsden (University of Waterloo, Waterloo, ON). Dr. M. Marsden purchased *X. laevis* from Nasco (Fort Atkinson, Wisconsin, USA) and these frogs were previously used in a breeding colony. Frogs were fed *ad libitum* with *Xenopus* food pellets (Boreal Science) and housed in opaque tanks filled with well water that was replaced twice a week. Frogs were kept at 22°C in the Aquatic Facility of the Biology building, University of Waterloo. Prior to tissue extraction, frogs were euthanized by immersion in a solution of benzocaine (300 mg/L). The frogs in the Aquatic Facility were maintained according to the guidelines of the University of Waterloo Animal Care Committee and the Canadian Council of Animal Care (CCAC-Canada).

### 2.2. Media

Leibovitz’s L-15 medium (HyClone) was diluted seven parts L-15 with three parts cell culture water (Lonza) to adjust for the osmolarity of amphibian cells and will herein be referred to as amphibian L-15 (AL-15). During initial establishment of cell tissue explant cultures, media was comprised of AL-15 supplemented with 20% fetal bovine serum (FBS; HyClone or VWR), 200 U/mL penicillin (HyClone), 200 μg/mL streptomycin (HyClone), 50 μg/mL gentamicin sulfate (HyClone) and 1 μg/mL amphotericin B (HyClone), herein referred to as establishment media. Once the primary cultures were established and in the early passage stage (passage 4-6), the continuous skin cell cultures were maintained using AL-15 medium supplemented with 15% FBS, 100 U/mL penicillin, 100 μg/mL streptomycin, and 50 μg/mL gentamicin sulfate. From passage 10 onwards, cells were maintained in AL-15 media supplemented with 15% FBS, without antibiotics, and will be referred to as complete medium. Amphibian phosphate-buffered saline (ADPBS) was prepared by diluting seven parts of Dulbecco’s phosphate-buffered saline (DPBS; Lonza) with 3 parts cell culture water. Similarly, seven parts 0.25% Trypsin-EDTA (HyClone) was combined with three parts cell culture water to adjust the osmolarity for use with amphibian cells and will be referred to as 0.175% Trypsin-EDTA.

### 2.3. Generation and establishment of continuous skin cultures

Dorsal and ventral skin tissues were dissected from euthanized *X. laevis* under sterile conditions and placed in individual petri dishes containing ADPBS supplemented with 200 U/mL penicillin, 200 μg/mL streptomycin, and 50 μg/mL gentamicin sulfate. Tissues were cut into small pieces with a scalpel and washed twice in ADPBS containing antibiotics prior to placing 4-6 tissue pieces into a 25 cm^2^ plug-seal tissue culture treated flask (BioLite, Thermo Scientific) containing 1.5 mL establishment medium to cover the tissue pieces. Primary tissue explant cultures were maintained at 26°C, in the absence of additional CO_2_. Medium was changed twice a week for the first three weeks using establishment media, and every 5-7 d afterwards using complete medium. Tissue culture explants were monitored for cell outgrowth. Once tissue explant cultures exhibited significant cell growth, adherent cells were collected by removing complete medium and washing cell cultures with 1× ADPBS prior to the addition of 0.175% Trypsin-EDTA. Upon cell detachment, complete medium was added, and gently pipetted to dissociate cell clumps prior to sub-culturing. Two continuous cell cultures were generated from these primary tissue explant cultures, named Xela DS2 and Xela VS2 for *X. laevis* dorsal and ventral skin, respectively.

### 2.4. Cell culture maintenance

Early Xela DS2 and Xela VS2 cell cultures were passed every 7 to 10 d at a 1:2 split. Once established (greater than passage 25 for Xela DS2, greater than passage 30 for Xela VS2), Xela DS2 and Xela VS2 cultures were passed every 3-4 d at a 1:4 split. Briefly, medium was removed from the flask and cells were washed with ADPBS. Following removal of the ADPBS, 0.175% trypsin-EDTA was added to detach cells from the cell culture vessel. Cells were subcultured at the indicated ratio into plug-seal tissue culture treated flasks containing fresh complete media and placed at 26°C in the absence of additional CO_2_. Cell morphology was monitored over successive passages by capturing phase-contrast images using a Nikon Eclipse TSX-100 microscope (taken at 100× magnification) fitted with a color camera and Picture Project software during early passages and with a Zeiss AxioVert-A1 microscope (taken at 200× magnification) fitted with an Axiocam 503 color camera and Zen 2.0 Lite software during later passages.

### 2.5. Plating efficiency

To determine plating efficiency, 40,000 cells per well were seeded in complete media and allowed to adhere overnight. The following day, adherent cells were collected via trypsin treatment and the number of viable cells were enumerated by mixing cells 1:1 with a solution of trypan blue (Invitrogen) and counting on a hemocytometer. Plating efficiencies were determined to be 79% for Xela DS2 and 83% for Xela VS2. For all experiments, initial seeding densities were adjusted for the individual cell lines to ensure that the desired number of adherent, viable cells per well was achieved. The reported cell numbers stated in the experiments described below represent the final number of cells that would be present in the tissue culture vessel the following day and accounted for differences in plating efficiencies for each cell line.

### 2.6. Cryopreservation

After harvest from tissue culture vessels, Xela DS2 or Xela VS2 cells were centrifuged at 500 × *g* for 5 min to pellet cells. The media was removed and cells were resuspended to approximately 2 × 10^6^ million cells per mL in chilled complete medium containing 10% DMSO (Fisher Scientific). One mL of cell suspension was aliquoted into individual cryogenic vials (Nunc) and vials were placed in a Mr. Frosty (Thermo Scientific) freezing container at −80°C overnight prior to long-term storage in liquid nitrogen.

### 2.7. Hematoxylin-Eosin (H&E) staining

Xela DS2 (passage 74) and Xela VS2 (passage 75) cells were seeded in 6-well plates (BioLite, Thermo Scientific) at cell densities of 300,000 cells/well in 1 mL of complete medium and allowed to adhere overnight at 26°C. The following day, media was removed from the wells and cells were fixed to the tissue culture plate by adding 70% ethanol for 2 min with gentle agitation. The ethanol was aspirated and residual ethanol was evaporated before cells were rinsed with 1 mL of Scott’s Tap Water (240 mM NaHCO_3_, 135 mM MgSO_4_, pH 8.0) for 30 s, with agitation. Cells were stained with Hematoxylin Gill 3 solution (VWR) for 5 min with agitation and washed with six exchanges of 1 mL Scott’s Tap Water. Cells were counterstained for 15 s with agitation in non-acidified 1% (w/v) Eosin Y (Fisher Scientific) in distilled water prior to rinsing cells with 1 mL of an increasing ethanol series (70%, 95%, and 100%). Cells were air dried prior to capture of bright field images using a Zeiss AxioVert-A1 microscope fitted with an Axiocam 503 color camera and Zen 2.0 Lite software.

### 2.8. Determination of senescence-associated beta-galactosidase activity

Cells were seeded in 12-well plates at cell densities of 50,000 cells/well for Xela DS2 (passage 9 and passage 74) and Xela VS2 (passage 13 and passage 75), or 30,000 cells/well for Xela BMW3 (passage 11 and passage 20) in 0.5 mL of complete medium. Cells were allowed to adhere overnight at 26°C prior to being assessed for β-galactosidase activity associated with cellular senescence using a commercially available Senescence Cells Histochemical Staining Kit (Sigma Aldrich). The manufacturer’s protocol was followed with the following modifications: plates were incubated at 26°C instead of 37°C after adding the staining mixture and washes were performed using ADPBS. Phase-contrast images were captured using a Nikon Eclipse TSX-100 at the earlier passage and a Zeiss AxioVert-A1 at the later passage. Digital images were taken of 3-5 fields of view, with greater than 400 cells counted, from each well for each cell type. The number of cells positive for senescence-associated β-galactosidase activity was expressed as a percent of the total number of cells enumerated for each cell type examined.

### 2.9. Hoechst staining for *Mycoplasma* sp. detection

Thirty thousand Xela DS2 (passage 151) and Xela VS2 (passage 135) cells were seeded separately onto sterile cover slips (Baxter) in a 6-well plate in 1 mL of complete medium. The plate was sealed with parafilm and placed overnight at 26°C. The following day, cells were fixed with an ice-cold fixative (1:3 v/v of acetic acid and methanol) for 20 min at – 20°C and incubated in the dark with 1 mL chilled Hoechst 33342 (Fisher) diluted 1:2000 in ADPBS prior to mounting on a glass slide in a 1:10 ratio of 100 mM Tris solution (pH 8.0) to glycerol. The cells were washed twice with room temperature ADPBS for 5 min with gentle agitation between each step. Images were captured on an inverted fluorescent microscope (Zeiss Axio Vert.A1) under oil immersion at 630× magnification.

### 2.10. Determination of optimal growth conditions

To assess the optimal growth conditions for Xela DS2 and Xela VS2, cells were seeded in 6-well plates at the desired concentration in 1 mL of complete media in duplicate or triplicate wells per time point and treatment and allowed to adhere at 26°C overnight. The following day, media was removed and replaced with 2 mL of the desired media and then placed at the indicated temperature. At each time point, cells were harvested by trypsin treatment and viable cells enumerated using trypan blue and a haemocytometer. In temperature experiments, Xela DS2 (passages 118-120) and Xela VS2 (between passages 123-129) were seeded at 40,000 cells/well and the plates shifted to temperatures of 14°C, 18°C, 22°C, 26°C, or 30°C. Cells were enumerated at 0 d, 2 d, 4 d, 6 d, and 8 d post transfer to the desired temperature. In FBS supplementation experiments, Xela DS2 (between passages 118-120) and Xela VS2 (between passages 123-129) were seeded at 40,000 cells/well and incubated with AL-15 supplemented with either 0%, 2%, 5%, 10%, 15%, or 20% FBS and all plates were incubated at 26°C. Cells were enumerated on 0 d, 4 d and 8 d post addition of FBS supplemented media. To examine the effect of cell density on cell growth, Xela DS2 (between passages 92-102) or Xela VS2 (between passages 93-103) cells were seeded at 10,000, 20,000 or 40,000 cells/well. Cells were incubated in complete medium at 26°C for the duration of the experiment and the number of viable cells enumerated at 0 d, 2 d, 4 d, 6 d, and 8 d. Temperature and FBS supplementation experiments were conducted three independent times and the cell density experiment was conducted five independent times.

### 2.11. DNA isolation

Total DNA was isolated from 2 × 10^6^ Xela DS2 or Xela VS2 cells using TRI reagent (Sigma) according to the manufacturer’s instructions. Briefly, cells were collected by centrifugation at 500 × *g* for 5 min and lysed with 1 mL of TRI reagent by pipetting. Samples were incubated at room temperature for 5 min prior to the addition of 0.2 mL chloroform and vigorous shaking for 15 s. Mixtures were allowed to sit at room temperature for 5 min prior to centrifugation at 12,000 × *g* for 15 min at 4°C. The top aqueous phase of each sample was discarded and 0.3 mL of 100% ethanol added to the remaining interphase and organic phase. The samples were mixed by inversion. Samples were left at room temperature for 3 min before centrifugation at 2,000 × *g* for 5 min at 4°C. The supernatants were removed and the DNA pellets were washed 2 × with 1 mL of 0.1 M sodium citrate in 10% ethanol. Between washes, the DNA pellets were allowed to stand in the sodium citrate/ethanol wash solution for 30 min, vortexed and centrifuged at 2,000 × *g* for 5 min at 4°C. The supernatants were removed and the DNA pellets were further washed 2 × with 1.5 mL 75% ethanol. Between washes, the DNA pellets were allowed to stand in a 75% ethanol solution for 20 min, vortexed and centrifuged at 2,000 × *g* for 5 min at 4°C. The ethanol was aspirated from each sample and the DNA pellets allowed to air dry for approximately 10 min before being resuspended in 1 × Tris-EDTA (TE; 500 μL pH 8.0 100 mM EDTA, 500 μl 1.0 M Tris-HCl up to 50 mL with DEPC-treated MilliQ water) buffer (pH 8.0). Total DNA quantity and purity of the extracted DNA samples were determined using a NanoDrop 1000 Spectrophotometer.

### 2.12. DNA barcoding

PCR reaction conditions and barcoding primers for amphibians were adapted from Che et al, 2012. Taxonomic DNA barcoding sequences were developed against the 5’-region of cytochrome c oxidase subunit I (*coi*) mitochondrial gene; for amphibians, the forward degenerate primer sequence was 5’-TYTCWACWAAYCAYAAAGAYATCGG-3’ and reverse degenerate primer sequence was 5’-ACYTCRGGRTGRCCRAARAATCA-3’ (Che et al., 2012). PCR reactions consisted of (final concentrations) PCR reaction buffer containing 2 mM MgCl_2_ (GeneDireX), 200 μM dNTP (GeneDireX), 200 μM sense and antisense primers (Sigma), 0.625 U *Taq* DNA Polymerase (GeneDireX), and 50 ng DNA template in a 25 μL reaction volume. The cycling conditions were as follows: initial denaturation at 95°C for 10 min; 35 amplification cycles of 94°C for 1 min, annealing at 45°C for 1 min and elongation at 72°C for 1 min; followed by a final extension step at 72°C for 10 min. In tandem with the PCR, a no template control using molecular grade water was used. Following amplification, 5 μL of 6× loading buffer [0.15% xylene cyanol (ICN Biomedicals), 30% v/v glycerol (EMD Chemicals)] was added to each reaction prior to loading into a 2.0% agarose (VWR) gel containing 1× RedSafe Nucleic Acid Staining Solution (FroggaBio) and electrophoresed in 1× Tris-Acetate-EDTA (TAE; from 50 × TAE, 242 g Tris-HCl, 100 mL pH 8.0 0.5 M EDTA, 57.1 mL glacial acetic acid up to 1.0 L with deionized water) buffer at 140 V for 20 min.

To purify DNA amplicons from agarose gel slices, bands were excised from the gel, placed in a 1.5 mL microcentrifuge tube and frozen at −80°C for 30 min. A hole was punctured in the bottom of a 0.5 mL tube (LifeGene), stuffed with a small amount of glass wool (Acros Organics), and the frozen gel slice added. The 0.5 mL tube containing the gel slice was placed inside a 1.5 mL microcentrifuge tube and centrifuged at 14,500 × *g* for 5 min. To the flow through, 0.1 volume of 3 M sodium acetate and 1 volume of isopropanol were added. Samples were vortexed before centrifugation at 14,500 × *g* for 15 min. The pellet was washed 2 × with 1 mL of 70% ethanol, vortexed and centrifuged at 14,500 × *g* for 5 min at 4 °C between washes. The precipitated DNA amplicons were resuspended in molecular grade water, cloned into the pUCM-T cloning vector (BioBasic) according to manufacturer’s instructions, and transformed into chemically competent *Escherichia coli* XL1-Blue by heat shock (1 min, 42°C), followed by recovery for 1 h at 37°C with shaking at 220 rpm in SOB medium (2% w/v tryptone, 0.5% w/v yeast extract, 8.56 mM NaCl, 2.5 mM KCl, 10 mM MgCl_2_). Transformed bacteria were plated on LB agar plates containing 100 μg/mL ampicillin (Fisher), 2 mg X-gal (BioBasic) and 2 μmol Isopropyl-β-D-thiogalactoside (IPTG; Fisher) at 37°C overnight. The following day, white colonies were selected and the presence of inserts confirmed by colony PCR using M13 forward (5’-TTGTAAAACCGACGGCCAGTG -3’) and reverse (5’-GGAAACAGCTATGACCATGATTACGC -3’) primers. Positive colonies were inoculated in 2 mL of LB broth containing 100 μg/mL ampicillin and grown overnight at 37°C with shaking at 220 rpm. The next day, bacteria were collected by centrifugation (10,000 × *g*, 5 min) and plasmids isolated using a modified plasmid isolation protocol. Bacterial pellets were resuspended in 100 μL of ice-cold 50 mM glucose, 10 mM EDTA, 25 mM Tris, pH 8.0 by vortexing. To this mixture, 200 μL of room temperature 0.2 N NaOH/1.0% SDS was added to the suspension and inverted to mix, followed by the addition of 150 μL of ice-cold 3 M KOAc, pH 6.0. Lysates were centrifuged at 15,500 × *g* for 30 min at 4°C. The supernatant was transferred to a new 1.5 mL microcentrifuge tube and incubated with 20 μg RNase A (BioBasic) for 20 min at 37°C. After incubation, 2.5 volumes of isopropanol was added, the solution inverted to mix, and centrifuged at 15,500 × *g* for 30 min at 4°C. The plasmid pellet was washed 2 × with 1 mL of 70% ethanol by vortexing and centrifuging at 15,500 × *g* for 5 min at 4°C between washes. Ethanol was aspirated and the plasmid pellet allowed to air dry before being resuspended in molecular grade water. Ten positive clones generated from each cell line were sent to the TCAG Facility at The Centre for Applied Genomics (Toronto, Ontario, Canada) for sequencing. Resulting sequences were trimmed to remove primer bases as well as any extended bases sequenced from the cloning vector. Processed sequences were subjected to *in silico* analysis via BLASTn (Altschul et al., 1990) against the *X. laevis* mitochondrial genome (GenBank Accession HM991335.1) to confirm the identity of the amplicons. Multiple sequence alignments of the Xela DS2 and Xela VS2 cytochrome c oxidase subunit 1 sequences were aligned against the *X. laevis* mitochondrial genome using Clustal Omega via EMBL-EBI (Madeira et al., 2019) and viewed using Jalview software (Waterhouse et al., 2009).

### 2.13. Total RNA isolation and cDNA synthesis

Total RNA was isolated from 1×10^6^ Xela DS2 or Xela VS2 cells using the EZ-10 Spin Column Total RNA Minipreps Super Kit (Bio Basic Canada Inc.) according to manufacturer’s instructions, with the following modification in order to include an on-column DNaseI digestion: Following the first RW wash step, 30 μL of DNase I solution containing 5 U of DNase I (Thermo Scientific) was added to the spin column for 20 min at room temperature. Afterwards, 500 μL of RW solution was added to the column and left at room temperature for 2-3 min before spin column centrifugation at 6,000 × *g* for 1 min. Isolation of total RNA from *X. laevis* skin tissues, previously stored at −80°C, was performed using TRI reagent according to the manufacturer’s instructions with the addition of a DNaseI (Fermentas) treatment according to the manufacturer specifications. Total RNA quantity and purity was determined using a NanoDrop 1000 Spectrophotometer. RNA quality was assessed by running 1 μg RNA on a 2.0% agarose gel containing 1× RedSafe Nucleic Acid Staining Solution and electrophoresed in 1× TAE buffer at 120 V for 30 min.

RNA was reverse transcribed into cDNA using the SensiFAST cDNA Synthesis Kit (BioLine) according to manufacturer’s specifications. Briefly, 500 ng of RNA was mixed with 4 μL of 5× reaction mix in a total volume of 20 μL. The resulting reactions were incubated at 25°C for 10 min, 42°C for 15 min and the reaction inactivated by incubation at 85°C for 5 min. Synthesized cDNA was stored at −20°C until use.

### 2.14. Reverse transcriptase polymerase chain reaction (RT-PCR) for the detection of cell type molecular markers

Primers for *X. laevis* β-actin (*actb*), collagen type I A1 (*col1a1*), collagen type I A2 (*col1a2*), collagen type III (*col3*), cytokeratin 5 (*krt5*), cytokeratin 14 (*krt14*), cytokeratin 19 (*krt19*), vimentin (*vim*), cadherin 1 (*cdh1*), claudin 1 (*cldn1*), claudin 3 (*cldn3*), and occludin (*ocln*) (Table 1) were designed using the online PrimerQuest tool from Integrated DNA Technologies (IDT; https://www.idtdna.com/Primerquest/Home/Index). RT-PCR reactions (25 μL reactions) consisted of (final concentrations) PCR reaction buffer containing 2 mM MgCl_2_, 200 μM dNTP, 200 μM sense and antisense primers (Sigma), 0.625 U Taq DNA polymerase and cDNA template. Reaction template included cDNA generated from isolated *X. laevis* dorsal skin and ventral skin tissues, and cDNA generated from Xela DS2 (between passages 72-88) and Xela VS2 (between passages 73-89). For cell type marker detection (*col1a1*, *col1a2*, *col3*, *krt5*, *krt14*, *krt19*, *vim*, *actb*) the cycling conditions were as follows: initial denaturation at 95°C for 5 min; 35 amplification cycles (*krt5*), 32 amplification cycles (*col1a1*, *col1a2*, *col3*, *krt14*, *krt19*), or 26 amplification cycles (*vim*, *actb*) of 95°C for 45 s, annealing at 55°C for 30 s and elongation at 72°C for 45 s; followed by a final extension step at 72°C for 5 min. For gap junction marker detection (*cdh1*, *cldn1*, *cldn3*, *ocln*, *actb*) the cycling conditions were as follows: initial denaturation at 95°C for 5 min; 28 amplification cycles (*cdh1*, *cldn1*, *cldn3*, *ocln*) or 23 amplification cycles *(actb)* of 95°C for 45 s, annealing at 57°C for 30 s and elongation at 72°C for 45 s; followed by a final extension step at 72°C for 5 min. A no template control was set up for all primer sets, and a RNA only control for each RNA sample was set up using β-actin primers. Both sets of RT-PCR reactions were repeated three times, using cDNA generated from Xela DS2 and Xela VS2 RNA at three different passages. To each RT-PCR reaction, 5 μL of 6× loading buffer (0.15% xylene cyanol, 30% v/v) was added prior to loading into a 2.0% agarose gel containing 1× RedSafe Nucleic Acid Staining Solution and electrophoresed in 1× TAE buffer at 140 V for 20 min. Gels were imaged via ChemiDoc imager using Image Lab program. Bands were excised from the gel, purified, and prepared for sequencing as described in Section 2.12. Resulting sequences were subjected to BLASTn (Altschul et al., 1990) analysis to confirm the identity of the amplicons.

**Table 1.**
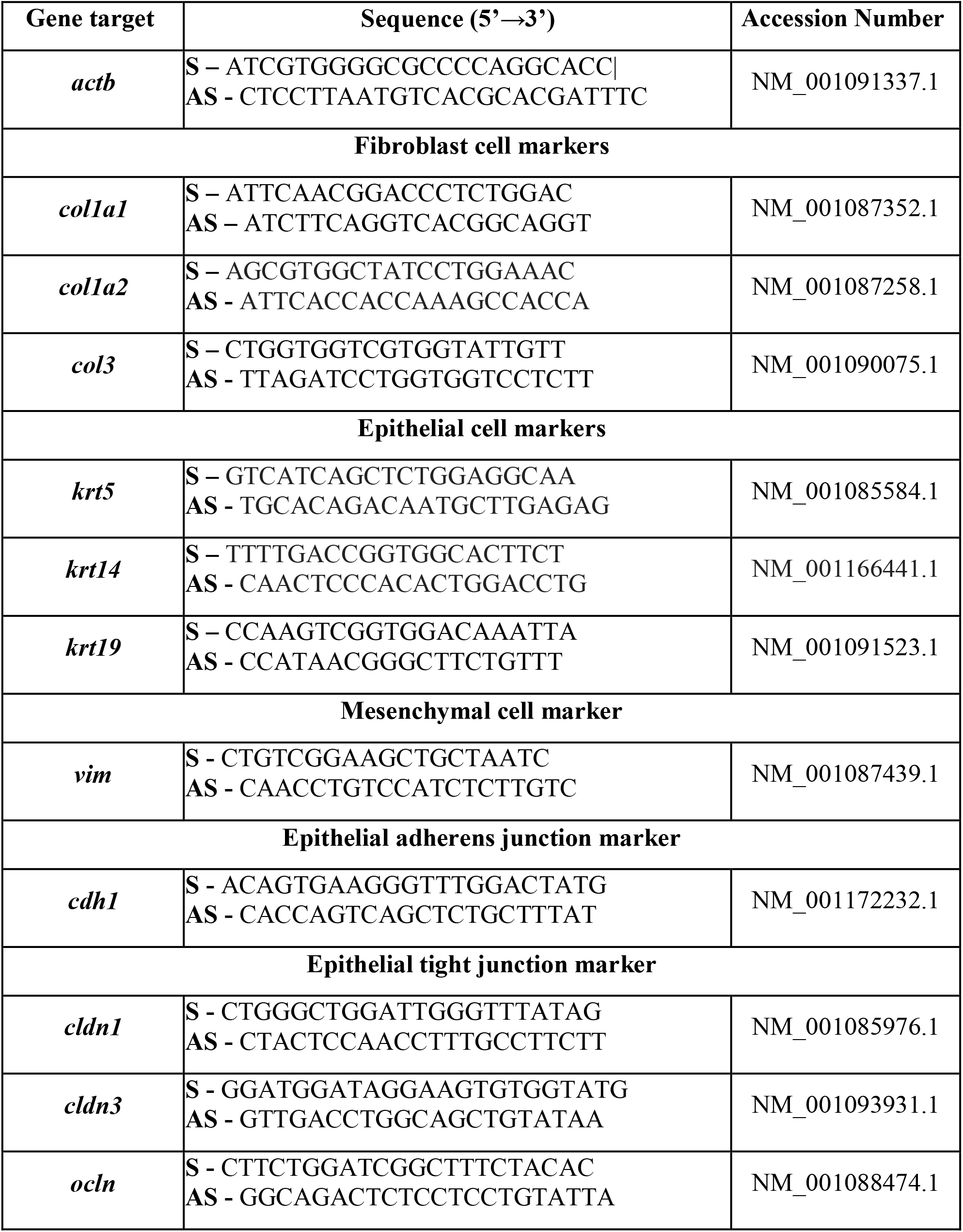
List of primers used for RT-PCR in the identification of Xela DS2 and Xela VS2 cell type.

### 2.15. Treatment of Xela DS2 and Xela VS2 cells with poly(I:C)

Two million Xela DS2 (between passages 110-116) or Xela VS2 (between passages 111-118) cells were seeded in a 6-well plate in 1 mL of complete medium and allowed to adhere overnight at 26°C. At the time of treatment, medium was replaced with 2 mL of fresh complete medium (time-matched controls) or complete medium containing 10 μg/mL polyinosinic-polycytidylic acid [poly(I:C), Sigma Aldrich Catalogue # P1530]. Phase contrast digital images were taken of control and poly(I:C) treated cells at 0 h, 3 h, 6 h, 12 h, and 24 h post treatment using a Leica DMi1 microscope fitted with a MC170 color camera and LASX 4.8 software to assess cell morphology. At each time point, cells from time-matched controls and poly(I:C) treatments were harvested by using a cell scraper to lift adherent cells from the culture vessel surface into suspension and the solution centrifuged at 230 × *g* for 10 min to collect both previously adherent cells as well as cells which lost adherence from poly(I:C) treatment. The supernatant was aspirated and the cell pellet was used for total RNA isolation using the EZ-10 Spin Column Total RNA Minipreps Super Kit as described in Section 2.13. Assessments of RNA quantity, purity, and integrity, and subsequent cDNA synthesis, were performed as described in Section 2.13. The cDNA samples were stored at −20°C until use in reverse transcriptase quantitative PCR (RT-qPCR) transcript analysis. This experiment was performed five independent times.

### 2.16. RT-qPCR of Xela DS2 and Xela VS2 poly(I:C) stimulated cells

Primer sequences and reference accession numbers for *X. laevis* tumor necrosis factor (*tnf*), interleukin-1 beta (*il1b*), interleukin-8 (*il8*), inhibitor of kappa B (*ikb*), type I interferon (*ifn1*), protein kinase R (*pkr*), myxovirus resistance gene 2 (*mx2*), *actb*, cyclophilin (*cyp*), elongation factor 1-α (*ef1a*), glyceraldehyde 3-phosphate dehydrogenase (*gapdh*), and hypoxanthine-guanine phosphoribosyltransferase (*hgprt*) can be found in Table 2; primer efficiencies and R^2^ values are also reported. Primer sequences for targets containing an asterisk (*) were accessed online in 2015 from Dr. Jacques Robert’s available primer list for *X. laevis* research (https://www.urmc.rochester.edu/microbiology-immunology/research/xenopus-laevis/primers.aspx). Prior to transcript analysis, gene stability testing was conducted for *actb*, *cyp*, *ef1a*, *gapdh*, and *hgprt* to determine a stable endogenous control selection. For gene stability testing, the transcript levels of each candidate reference gene was determined using cDNA samples from each treatment and time point generated from Section 2.15 RT-qPCR reactions were prepared in triplicate and consisted of (final concentrations) 250 nM sense and antisense primer, 5 μL PowerUp SYBR green mix (Thermo Scientific), and 2.5 μL of 1:40 diluted cDNA at a final reaction volume of 10 μL. All reactions were prepared in a MicroAmp fast optical 96-well reaction plate and optical film (Life Technologies) and run on QuantStudio5 Real-Time PCR System (Thermo Scientific). The cycling conditions were as follows: initial denaturation at 50°C for 2 min, followed by 95°C for 2 min, 40 amplification cycles of denaturation at 95°C for 1 s, and extension at 60°C for 30 s. A melt curve step followed all runs to ensure only a single dissociation peak was present, with initial denaturation at 95°C for 1 s, then dissociation analysis at 60°C for 20 s followed by 0.1°C/s increments up to 95°C. Gene stability measure (M-score) was determined for all endogenous control candidates (Suppl. Table 1), wherein a lower M-score infers stronger gene stability across time and treatment. The lowest M-score was for *ef1a* at 0.278 and 0.217 for Xela DS2 and Xela VS2 respectively (Suppl. Table 1); *ef1a* was thus selected as the endogenous control.

**Table 2.**
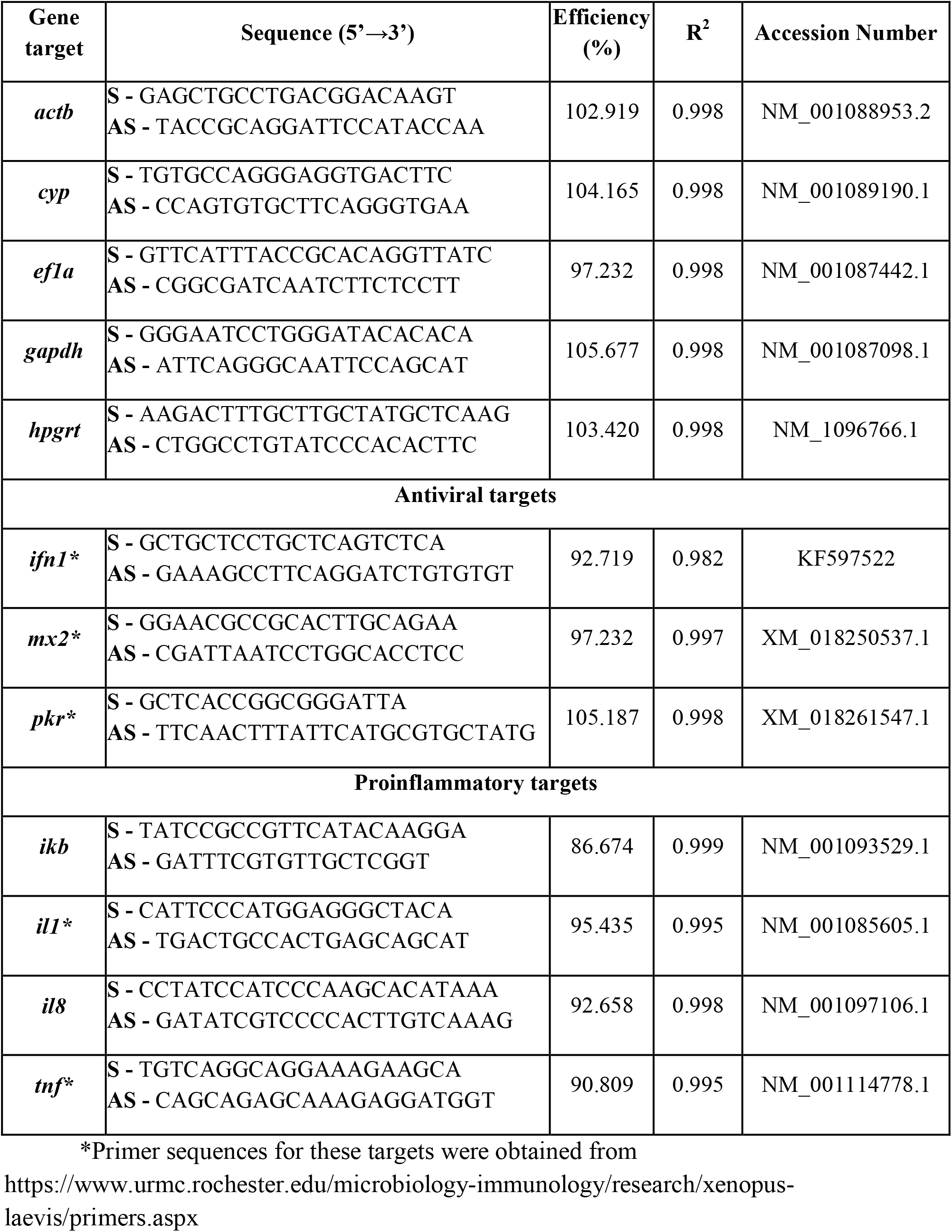
List of primers and efficiencies used for RT-qPCR in this study.

For transcript analysis, cDNA generated from Section 2.15 was diluted to 1:20 for all reactions. RT-qPCR reactions were prepared in triplicate and consisted of (final concentrations) 250 nM sense and antisense primer, 5 μL PowerUp SYBR green mix, and 2.5 μL of 1:20 diluted cDNA at a final reaction volume of 10 μL. The cycling conditions and melt curve step were performed as previously described. RT-qPCR reactions for transcript analysis was conducted using cDNA generated from Xela DS2 and Xela VS2 RNA at five different passages (as mentioned in Section 2.15; *N* = 5). Amplicons from all primer sets used in RT-qPCR reactions were sequenced as described in Section 2.12 and resulting sequences were subjected to BLASTn (Altschul et al., 1990) analysis to confirm the identity of the amplicons and amplification of a single gene target.

### 2.17. Statistics

For cell growth experiments, data were analyzed by a two-way analysis of variance (ANOVA) for the relationship between treatment and time and followed with a Tukey’s post hoc analysis. Groups were considered statistically significant groups when *p* < 0.05. For RT-qPCR analysis of Xela DS2 and Xela VS2 poly(I:C) stimulated cells, statistical analysis was performed by using a Mann-Whitney test with Bonferroni correction, which considers each time-point (4 total) as an independent test, correcting initial *p* < 0.05 to *p* < 0.0125. As such, statistical significance was determined within each time-point between treated cells and their time matched control wherein significant groups have *p* < 0.0125. Statistical analyses were performed using GraphPad Prism v6 software.

## 3. Results

### 3.1. Development of Xela DS2 and Xela VS2 cell lines and cell characteristics

In order to elucidate the contribution of skin epithelial cells to amphibian innate immune responses, we established skin epithelial cell lines from the dorsal and ventral skin of adult female *X. laevis*. Within one week of placing *X. laevis* dorsal skin or ventral skin tissue explants in tissue culture flasks, epithelial-like cells could be seen migrating out of the tissues (not shown). These cells appeared to have an epithelial-like morphology with simple epithelium structure as seen by a single layer of polygonal cells with close cell-to-cell contact. Over a period of 3 – 4 months, a flask containing dorsal skin fragments and a flask containing ventral skin fragments, originating from the same animal, produced large islands of adherent single-layered cells could be observed surrounding the central mass of tissue and covered ~60-70% of the flask. Trypsin treatment was used to readily detach the cells over subsequent passages, leading to continuous propagation. The resulting cell lines originating from *X. laevis* dorsal skin and ventral skin tissue explants were named Xela DS2 and Xela VS2, respectively. During this time, Xela DS2 and Xela VS2 consistently displayed epithelial-like morphology. At early cell passages of passage 9 for Xela DS2 (Fig. 1A) and passage 13 for Xela VS2 (Fig. 1E), cells were largely adherent and epithelial-like in morphology (Fig. 1A, E). While earlier passages of Xela DS2 and Xela VS2 had slow and patchy growth, later passages of Xela DS2 (Fig. 1B, C) and Xela VS2 (Fig. 1F, G) grew more rapidly and formed confluent monolayers. If cell cultures became over-confluent, cells would begin to lose adherence. Both Xela DS2 and Xela VS2 cell lines are now routinely maintained in complete medium without antibiotics and are split 1:4 every 3-4 days. Xela DS2 and Xela VS2 have been passed >160 times over 4 years since their initial establishment and have been successfully cryopreserved at a variety of passages in complete medium containing 10% DMSO. Xela DS2 and Xela VS2 cell lines were confirmed to be of *X. laevis* origin by sequencing a region of the cytochrome c oxidase I gene in the mitochondrial genome and comparison to the published *X. laevis* cytochrome c oxidase subunit I in the mitochondrial genome (GenBank Accession HM991335.1). Both Xela DS2 (Suppl. Fig. 1) and Xela VS2 (Suppl. Fig. 2) sequenced COI regions were a match to that of the published mitochondrial COI gene from *X. laevis*.

**Figure 1.**
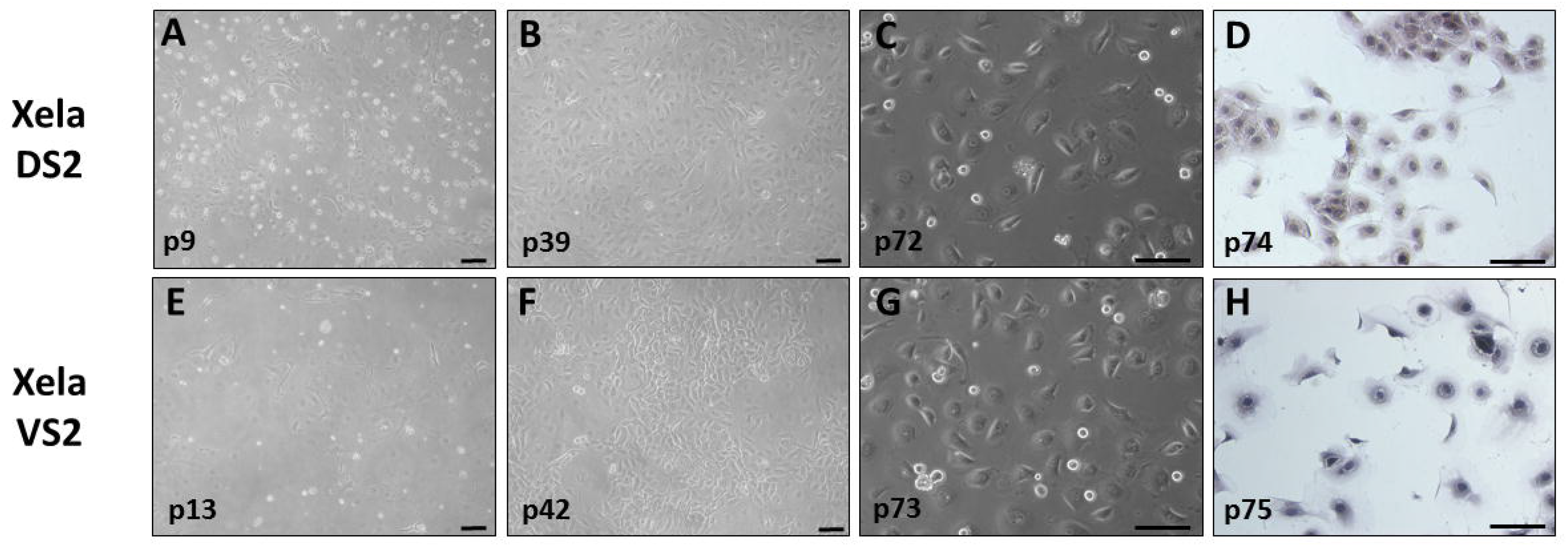
Xela DS2 and Xela VS2 cells exhibit epithelial cell-like morphology. Phase-contrast digital images of Xela DS2 cells were taken at passage 9 (A), passage 39 (B), and passage 72 (C), and of Xela VS2 cells at passage 13 (E), passage 42 (F), and passage 72 (G). Bright-field digital images of Hematoxylin and Eosin stained Xela DS2 cells (D) and Xela VS2 cells (H). Scale bars represent a distance of 100 μm.

To characterize cell morphology, cells were stained with Hematoxylin-Eosin. Xela DS2 cells were characterized by oval nuclei and light pink cytoplasm (Fig. 1D), while Xela VS2 cells also had oval nuclei but relatively clear to light purple cytoplasm (Fig. 1H) and both cell lines appeared epithelial-like with polygonal morphology. At lower cell densities, some Xela DS2 and Xela VS2 cells also appeared crescent-like (Fig. 1D, H). At higher cell densities, Xela DS2 and Xela VS2 become compact and had an average cell diameter of approximately 35 μm for both. At early passages, Xela DS2 (passage 9) and Xela VS2 (passage 13) exhibited 3% and 24% cellular senescence, respectively (Fig. 2C, E). However, with successive passages both Xela DS2 (Fig. 2D) and Xela VS2 (Fig. 2F) cells lost β-galactosidase activity. Cells from Xela BMW3 continuous cell cultures derived from *X. laevis* bone marrow were used as a positive control as these cells previously stained positive for β-galactosidase activity (unpublished observations). At passage 11 (Fig. 2A) and passage 20 (Fig. 2D), 16% and 46% of the BMW3 cells appeared blue, respectively, indicative of β-galactosidase activity. In addition, Xela DS2 and Xela VS2 were tested for potential mycoplasma contamination by employing the Hoechst staining method. Extranuclear fluorescent foci were not observed in Xela DS2 (Fig. 3A) or Xela VS2 (Fig. 3B) cultures and indicate these cultures are mycoplasma free.

**Figure 2.**
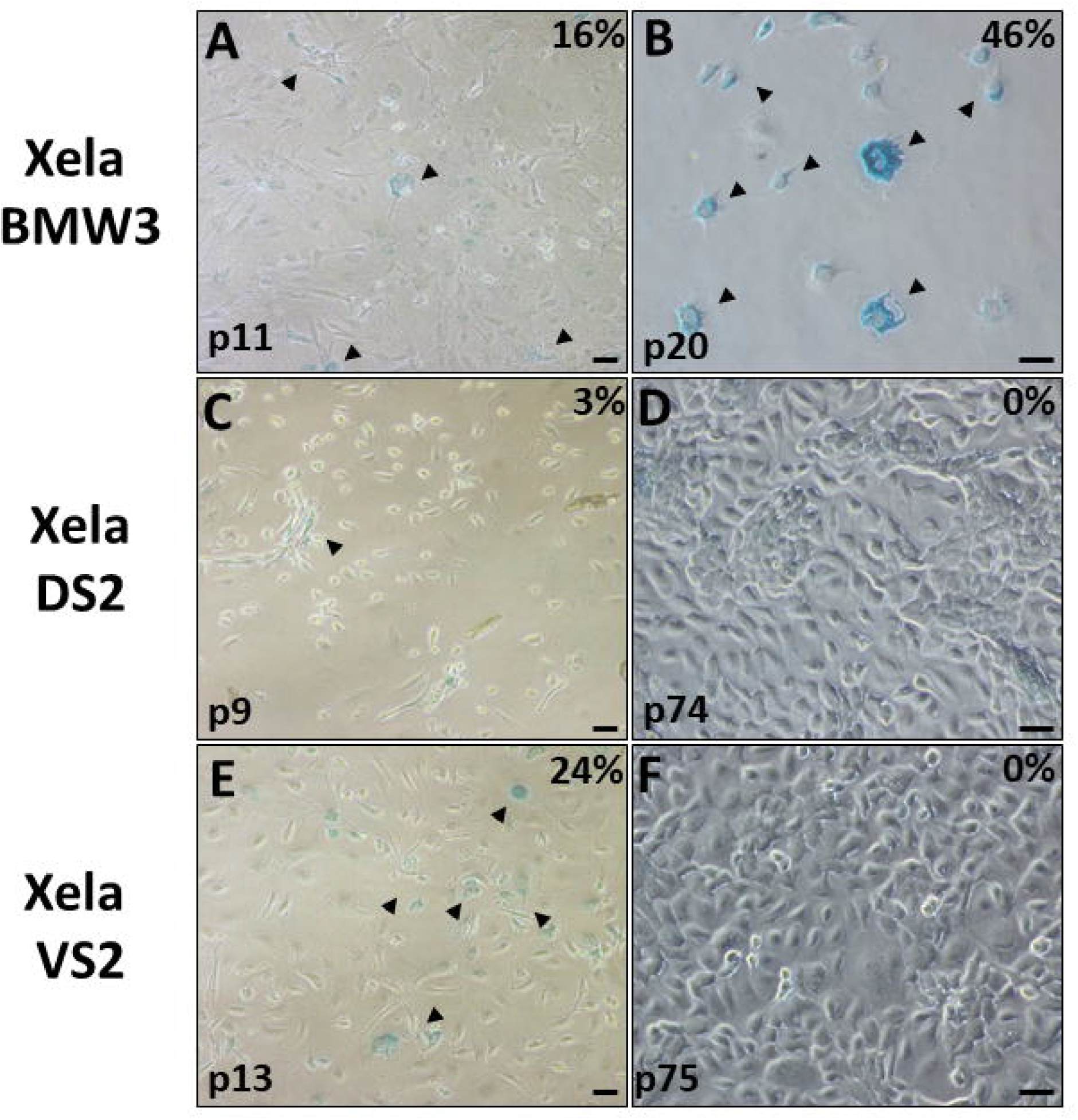
Senescence associated β-galactosidase activity of Xela DS2 and Xela VS2 cells from early and established cultures. Xela DS2 and Xela VS2 cells were tested for senescence associated β-galactosidase activity appear blue. Photomicrographs of Xela BMW3 cells at passage 11 (A), Xela DS2 cells at passage 9 (C), Xela VS2 cells at passage 13 (E), Xela BMW3 cells at passage 20 (B), Xela DS2 cells at passage 74 (D), and Xela VS2 cells at passage 75 (F). Xela BMW3 cells were used as a positive control for β-galactosidase activity. The percentages of β-galactosidase positive cells are indicated in the top-right hand corner of each image. Scale bars represent a distance of 50 μM.

**Figure 3.**
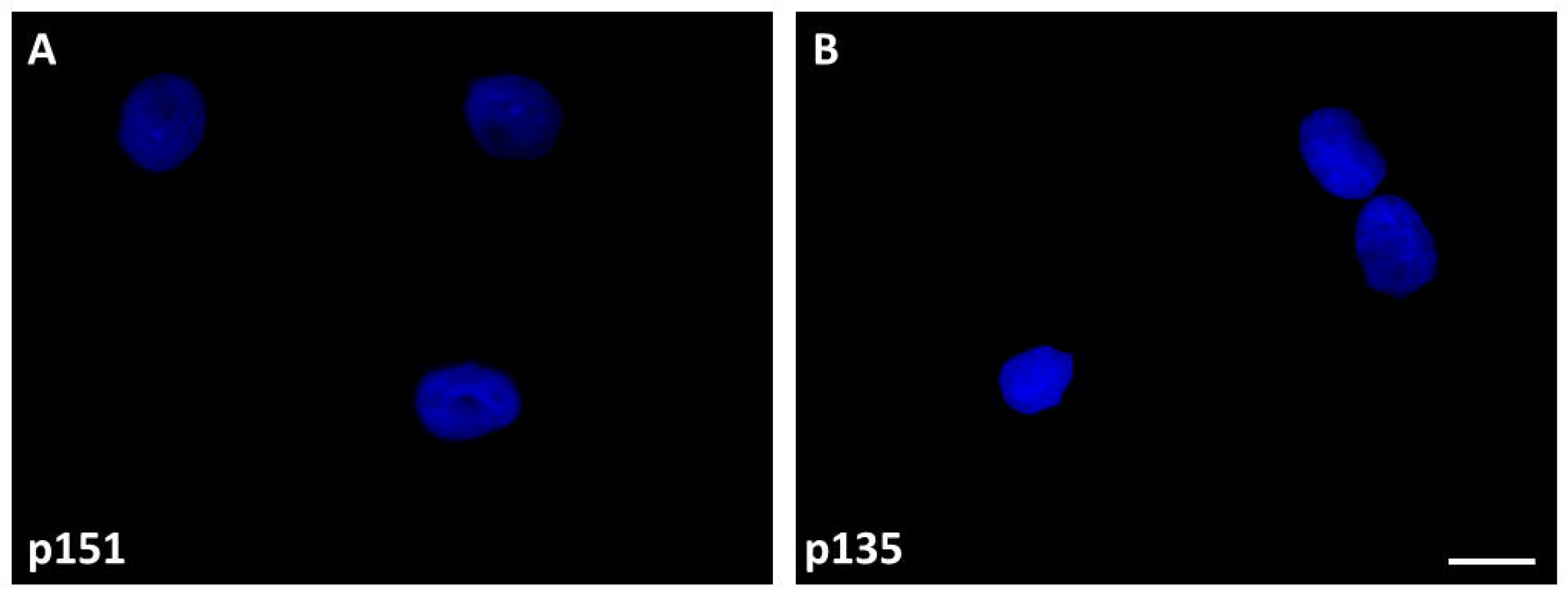
Hoechst staining of Xela DS2 and Xela VS2 cells. (A) Xela DS2 at passage 151 (B) Xela VS2 at passage 135. Both cell lines exhibit nuclei with no evidence of fluorescent specks or puncta outside of the nucleus. Magnification, 630× and scale bar, 20 μm.

### 3.2. Effect of culture conditions on Xela DS2 and Xela VS2 cell proliferation

Although other *X. laevis* cell lines are routinely cultivated in osmotically adjusted medium containing FBS, the percentage of FBS and incubation temperature varies with the cell line (Pudney et al., 1973; Rafferty, 1969). Therefore, we undertook studies to determine the optimal level of serum supplementation and incubation temperature for Xela DS2 and Xela VS2. Two-way ANOVA analysis indicated that there is a significant effect due to FBS treatment (p < 0.0001) and time (p < 0.0001) on Xela DS2 and Xela VS2 cell proliferation. In the absence of FBS supplementation, or with only 2% FBS, no significant differences in cell numbers were observed in Xela DS2 (Fig. 4A) or Xela VS2 (Fig. 4B) cultures over 8 days at 26°C. However, we noted that Xela DS2 and Xela VS2 cells started to lose adherence to the tissue culture vessel over time when cultured in the absence of FBS. Supplementation of cultures with 5% FBS also did not support statistically significant proliferation of Xela DS2 (Fig. 4A) or Xela VS2 cells (Fig. 4B), although there were roughly 4-fold as many cells by day 8 of culture for both cell lines. A minimum of 10% FBS was required to promote significant increases in cell numbers in Xela DS2 (Fig. 4A) or Xela VS2 (Fig. 4B) over 8 days. Indeed, fold change in Xela DS2 (Fig. 4A) and Xela VS2 (Fig. 4B) cell numbers increased in a dose-dependent manner with FBS supplementation after 8 days, achieving a maximum ~10-fold increase in cell numbers when cultured in the presence of a minimum of 15% FBS for Xela DS2 (Fig. 4A) or 10% FBS for Xela VS2 (Fig. 4B). Based on these results, 15% FBS supplementation of AL-15 media was chosen for optimal maintenance of Xela DS2 and Xela VS2.

**Figure 4.**
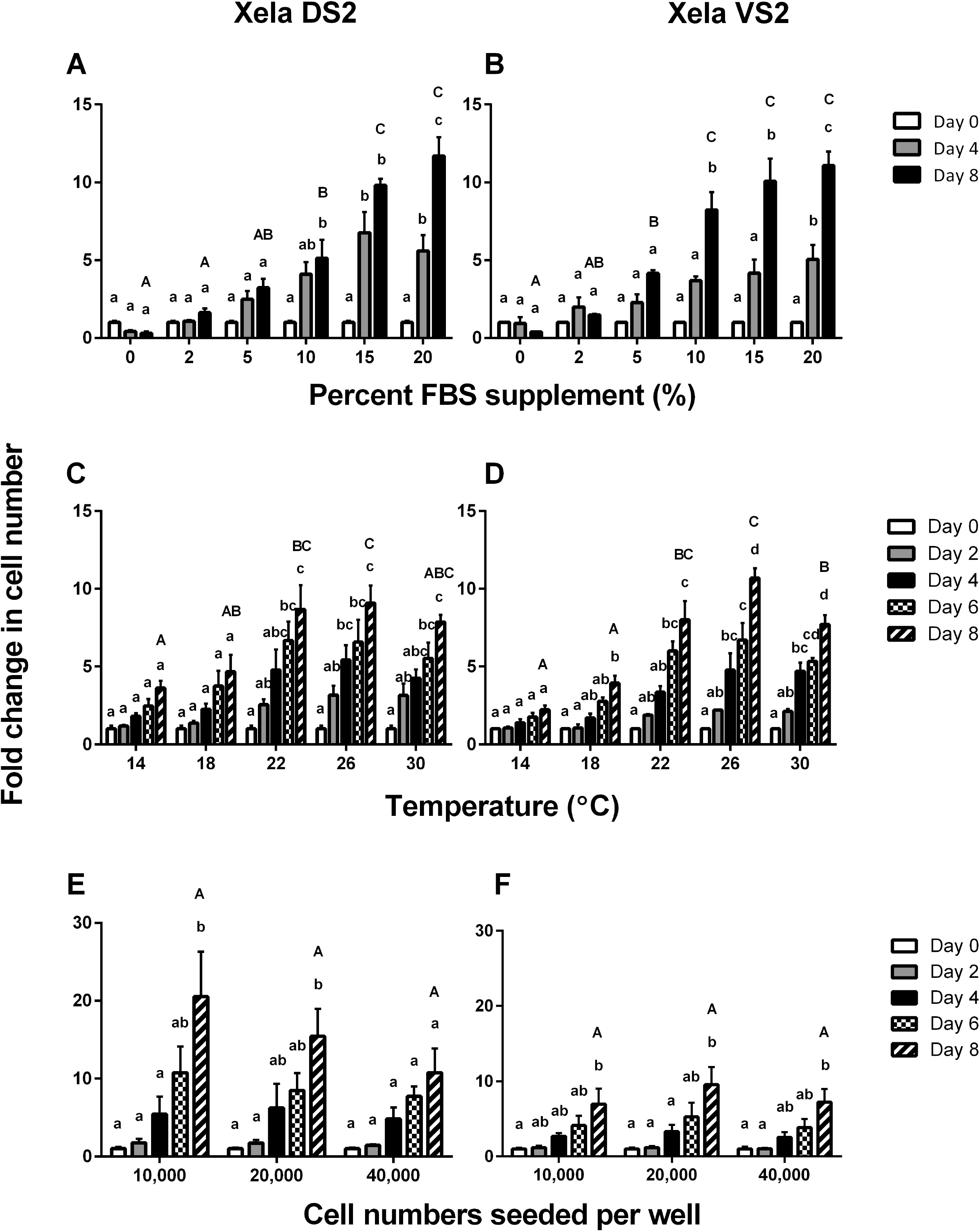
Fold change in cell growth for Xela DS2 and Xela VS2 cells under varying conditions. Fold change in Xela DS2 (A) and Xela VS2 (B) cell numbers when seeded at 80,000 cells per well and grown at 26 °C in AL-15 medium supplemented with 0%, 2%, 5%, 10%, 15%, 20%, or 30% FBS (n = 3 independent trials). Fold change in Xela DS2 (C) and Xela VS2 (D) cell numbers when seeded at 40,000 cells per well and grown in AL-15 supplemented with 20% FBS at varying temperatures (n = 3 independent trials). Fold change in Xela DS2 (E) and Xela VS2 (F) cell numbers after being seeded at varying initial cell densities and grown at 26 °C in AL-15 medium supplemented with 15% FBS (n = 5 independent trials). For all experiments, viable cells were enumerated on the indicated day and expressed as a fold change in cell number relative to cell numbers at the Day 0 time point; Day 0 counts were the same for all treatments within an experiment. Significant differences were determined by a two-way ANOVA and Tukey’s post-hoc test (*p* < 0.05), wherein like lettering indicates no statistical significance between groups.

A significant effect of incubation temperature (p = 0.01850 for Xela DS2, p < 0.0001 for Xela VS2) and time (p < 0.0001) on Xela DS2 and Xela VS2 proliferation was observed. Minimal proliferation of Xela DS2 (Fig. 4C) and Xela VS2 (Fig. 4D) was observed at lower temperatures of 14°C and 18°C after 8 d of cultivation. However, both Xela DS2 (Fig. 4C) and Xela VS2 (Fig. 4D) demonstrated significant proliferation over 8 d at temperatures above 22°C, with no significant difference in fold change in cell number at 8 days post seeding across when incubated at 22°C, 26°C and 30°C for Xela DS2 (Fig. 4C) or 22°C and 26°C for Xela VS2 (Fig. 4D). Over 8 d of cultivation, Xela DS2 cells exhibited a 9-fold, 9-fold and 8-fold change in cell number when grown at 22°C, 26°C and 30°C (Fig. 4C) respectively, while Xela VS2 exhibited an 8-fold, 10.5-fold, and 7.5-fold change in cell number at identical growth temperatures (Fig. 4D). Since Xela DS2 and Xela VS2 appeared to have optimal cell growth over 8 days of cultivation at 26°C, this temperature was selected to maintain the cells and to conduct further studies.

To assess if Xela DS2 or Xela VS2 cells demonstrated contact inhibition, cells were seeded at 10,000 cells/well, 20,000 cells/well, or 40,000 cells/well in complete medium and cultured for 8 days at 26°C. Two-way ANOVA analysis indicated that there is a significant effect due to time (p < 0.0001) but not seeding cell density (p = 0.5607 for Xela DS2, p = 0.6700 for Xela VS2) on Xela DS2 (Fig. 4E) and Xela VS2 (Fig. 4F) cell numbers.

### 3.3. Xela DS2 and Xela VS2 express molecular markers of epithelial cells

To characterize the cell type present in Xela DS2 and Xela VS2, the molecular signatures of Xela DS2 and Xela VS2 were investigated by selecting gene targets based on those typically expressed in mammalian fibroblast cells, epithelial cells and mesenchymal cells. Genes known to be expressed primarily in fibroblasts include *col1a1*, *col1a2*, and *col3* and are necessary for extracellular matrix formation (Miskulin et al., 1986). Epithelial cells express an array of cytokeratin for structural support such as *krt5* and *krt14* in epidermal stratified squamous epithelium, or *krt19* in simple epithelium (Chu and Weiss, 2002; Moll et al., 1982). Finally, *vim* was selected as a marker of mesenchymal cells since epithelial cells are known to undergo an epithelial to mesenchymal transition (EMT) in instances (Chaw et al., 2012). Dorsal and ventral skin tissues from *X. laevis* expressed all the gene targets examined (Fig. 5), with the exception of *krt14* which did not appear to be expressed in dorsal skin tissue. Xela DS2 demonstrated robust expressions of *col1a2*, *krt19*, and *vim*, and barely detectable expressions of *col1a1* and *col3* (Fig. 5). Expressions of *krt5* and *krt14* in Xela DS2 were not detected (Fig. 5). Xela VS2 expressed *col1a1*, *col1a2*, *col3*, *krt5*, *krt19*, and *vim* genes (Fig. 5). However, Xela VS2 did not appear to express *krt14* (Fig. 5).

**Figure 5.**
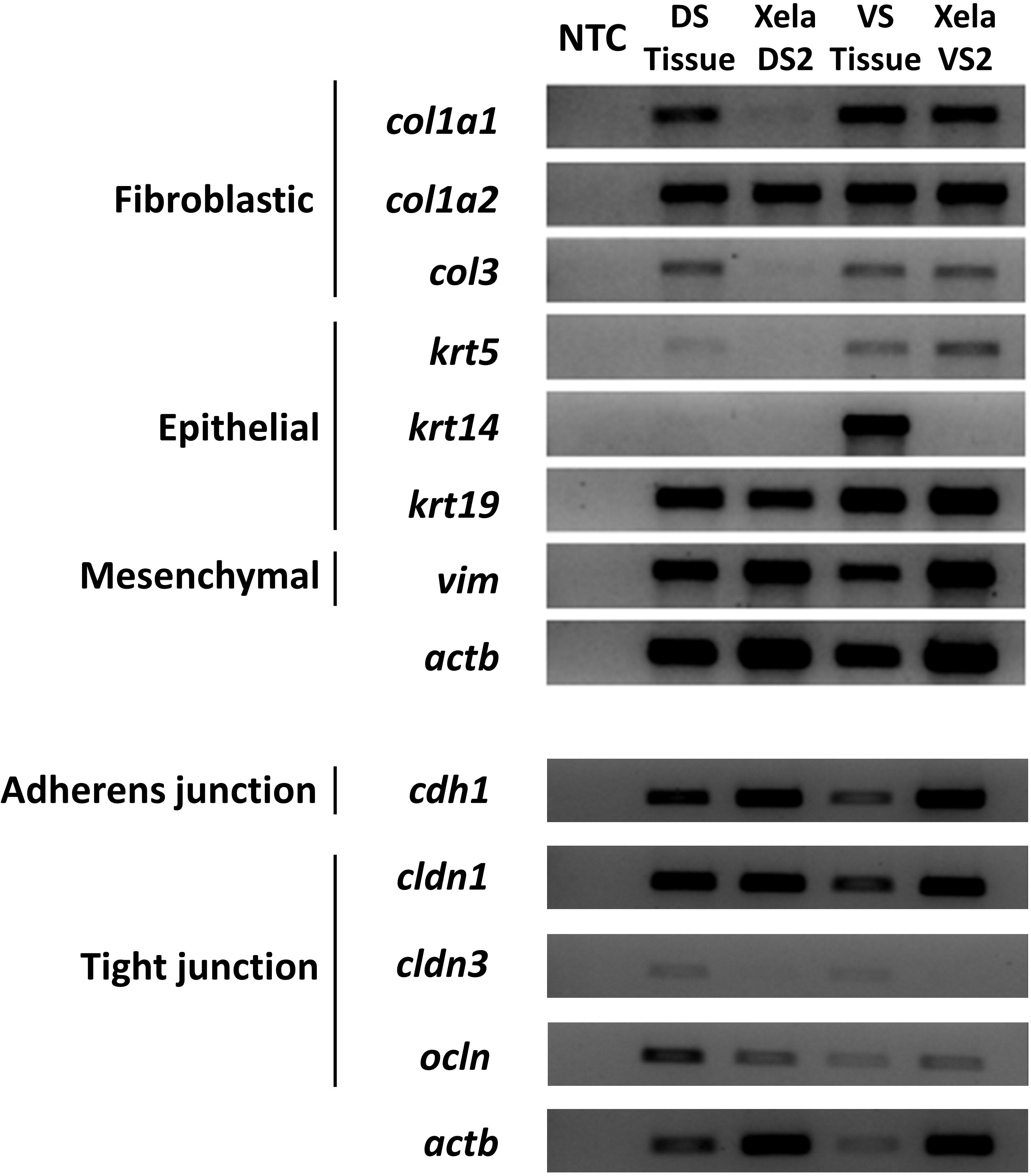
Xela DS2 and Xela VS2 have unique molecular signatures characteristic of epithelial cells. Gel electrophoresis from RT-PCR for cell type markers of dorsal skin tissue (DS Tissue), Xela DS2, ventral skin tissue (VS Tissue), and Xela VS2. Gene targets include fibroblastic molecular markers *col1a1*, *col1a2*, *col3*, epithelial molecular markers *krt5*, *krt14*, *krt19*, mesenchymal marker *vim*, and *actb* as an endogenous control. Gel electrophoresis from RT-PCR for epithelial gap junction markers was performed on the same set of samples. Gene targets include adherens junction marker *cdh1*, tight junction markers *cldn1*, *cldn3*, *ocln*, and *actb* as an endogenous control. NTC, no template control (n = 3 independent trials, representative trial shown).

To further ascertain the cell type identity of Xela DS2 and Xela VS2, the expressions of genes involved in encoding for tight and adherens junction proteins, essential to the integrity of the epithelial barrier (Menco, 1980; Villaro et al., 1998), were examined. Tight junctions are localized to epithelial and endothelial cells, requiring claudin and occludin family membrane proteins for formation, while adherens junctions require cadherin family proteins to be functional (Coyne et al., 2003; Yap et al., 1997). Both Xela DS2 and Xela VS2 expressed *cdh1*, *cldn1* and *ocln* genes (Fig. 5), wherein *cdh1* is strictly localized to epithelial cells in mammalian systems. The expression of *cldn3* is detected at relatively low levels in frog skin tissue but remains undetected in Xela DS2 or Xela VS2 (Fig. 5).

### 3.4. Effect of poly(I:C) treatment on Xela DS2 and Xela VS2 cell adherence

To assess the potential effects of viral dsRNA on Xela DS2 and Xela VS2, cells were treated with 10 μg/mL poly(I:C), a synthetic viral dsRNA analogue. A proportion of poly(I:C) treated cells exhibited a loss of adherence to the tissue culture vessel, appearing as suspension cells in the media within 3 h post treatment (Fig. 6). Following this initial loss of cell adherence, no further loss of cell attachment was observed over the 24 h treatment period (Fig. 6). Sampling of the suspension cells at the 24 h time point revealed the exclusion of trypan blue by these suspension cells, suggesting their membranes were intact and the cells viable at this time point. Non-treated Xela DS2 and Xela VS2 cells did not exhibit any changes in cell morphology or adherence to the tissue culture vessel over the 24 h time period (Fig. 6).

**Figure 6.**
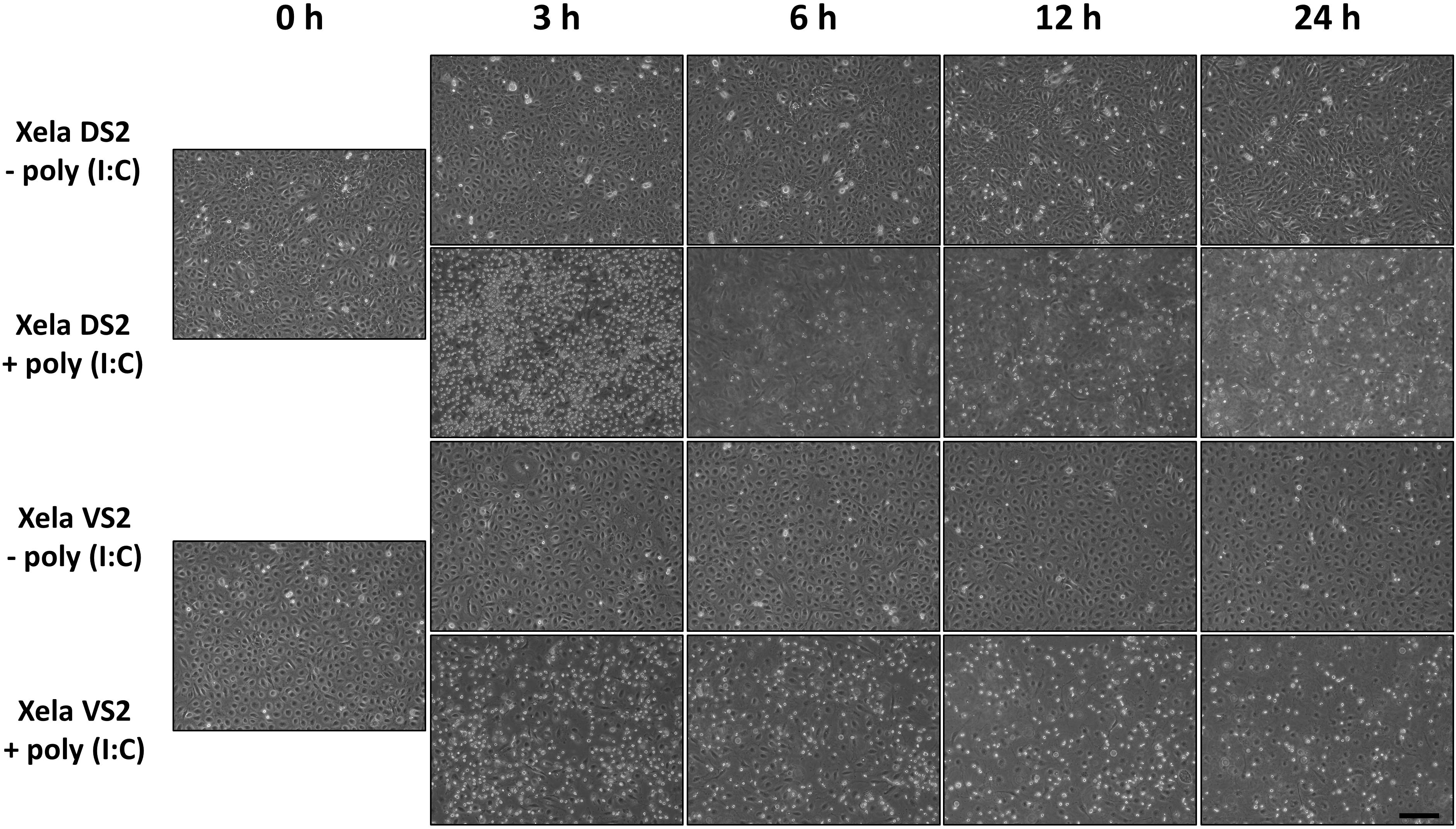
Xela DS2 and Xela VS2 cell morphology following poly(I:C) treatment. Phase contrast images showing the response of Xela DS2 and Xela VS2 cells to the absence or presence of 10 μg/mL of poly(I:C). Non-treated Xela DS2 or Xela VS2 (left panels) and poly(I:C) treated cells (right panels). Images were taken at 100× magnification and scale bar represents a distance of 200 μm.

### 3.5. Effect of poly(I:C) treatment on antiviral gene expressions in Xela DS2 and Xela VS2

We next assessed whether poly(I:C) might be a potent inducer of type I interferon (*ifn1*) in Xela DS2 and Xela VS2. As this set of experiments was conducted prior to the identification of the expanded *X. laevis* intron-containing and intronless IFNs (Sang et al., 2016), we used the *X. laevis ifn1* primers reported in (Grayfer et al., 2014). Subsequent *in silico* analysis suggests that the *ifn1* primer set used in this study is likely targeting the newly identified *ifn6* (Suppl. Fig. 3) within the expanded *X. laevis* Type I IFN family. Xela DS2 showed an initial 3.9 ± 0.7 fold (p = 0.0079) increase in *ifn1* transcripts at 3 h post treatment and transcript levels continued to increase in a time-dependent manner, reaching a 7.7 ± 2.6 fold (p = 0.0079) significant increase at 12 h post treatment relative to the time-matched control (Fig. 7A). Although *ifn1* transcript levels were 20.0 ± 9.7 fold (p = 0.0317) higher in Xela DS2 than the time matched non-treated control cells at 24 h post treatment, the transcript levels were not statistically significant. Xela VS2 also demonstrated a time-dependent increase in *ifn1* transcript levels over 24 h post poly(I:C) treatment (Fig. 7B). At 6 h post poly(I:C) treatment, *ifn1* transcript levels were 8.9 ± 6.3 fold that of the time-matched, non-treated control cells (Fig. 7B), although not statistically significant (p = 0.7937). By 12 h and 24 h post poly(I:C) treatment, *ifn-1* mRNA levels in Xela VS2 were 14.6 ± 8.4 fold (p = 0.0079) and 27.0 ± 14.8 fold higher (p =0.0159), respectively, than time-matched, non-treated controls (Fig. 7B). Although the 24 h *ifn-1* levels are not statistically significant, the p-value (p = 0.0159) is close to the cut-off (p < 0.0125).

**Figure 7.**
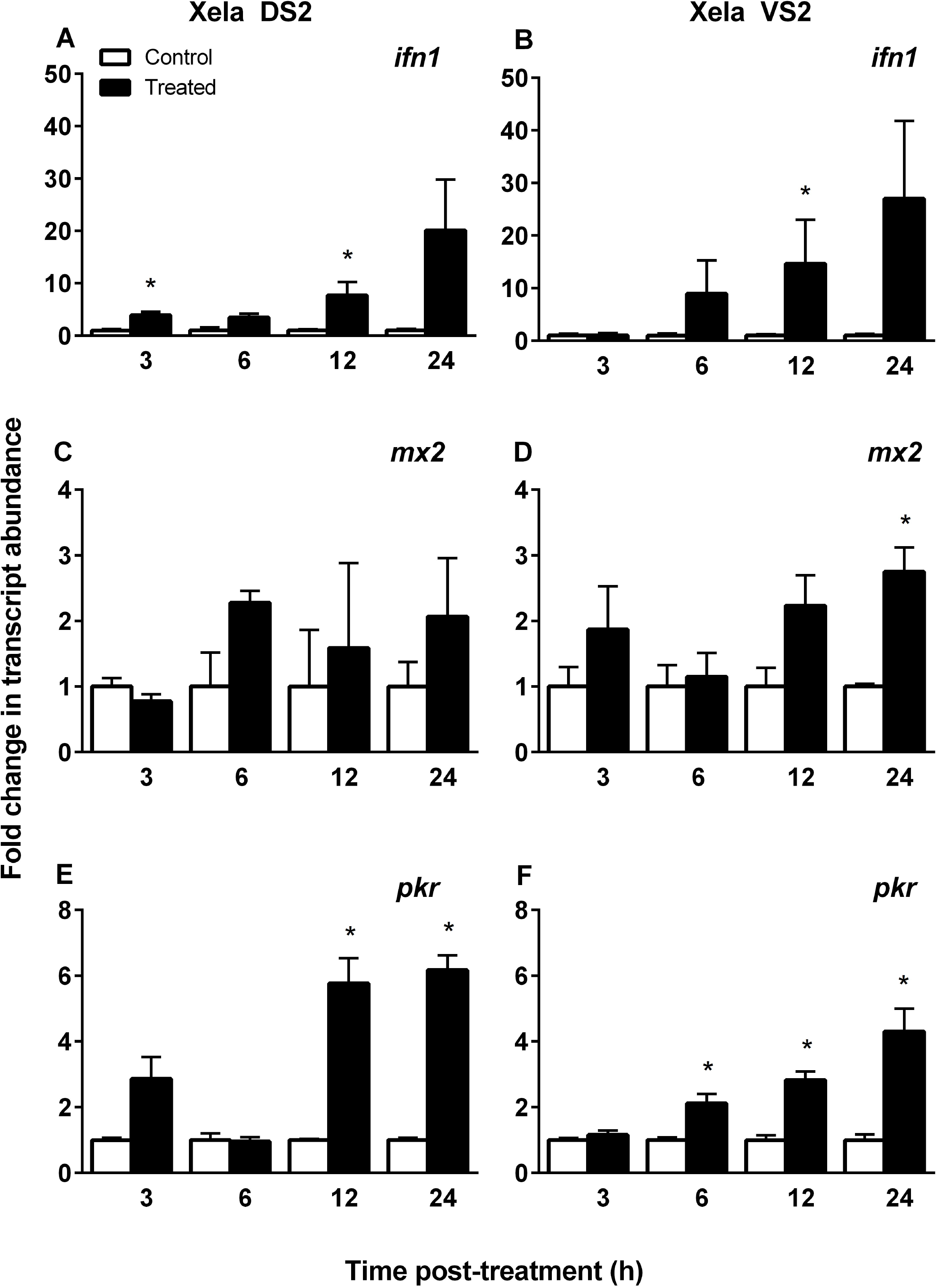
Poly(I:C) treatment induces upregulation of type I interferon and interferon stimulated genes in Xela DS2 and Xela VS2. RT-qPCR was performed on cDNA generated from non-treated (control) and poly(I:C) treated Xela DS2 or Xela VS2 cells to determine relative transcript levels of *ifn1* (A, B), *mx2* (C, D), and *pkr*. For all experiments, transcript levels from treated samples were normalized to that of the respective time matched controls, (n = 5 independent trials). Significant differences were determined by a Mann-Whitney test with Bonferroni correction (*p* < 0.0125).

Due to the unavailability of an antibody to detect the production and secretion of type I interferons in *X. laevis*, we assessed whether poly(I:C) treated Xela DS2 and Xela VS2 were upregulating the expressions of key downstream ISGs, namely *mx2* and *pkr*. Over 24 h post treatment with poly(I:C), *mx2* transcript levels did not significantly change in Xela DS2 (Fig. 7C) or Xela VS2, with the exception of the 24 h post treatment time point in Xela VS2 (2.8 ± 0.4 fold increase, p = 0.0079) (Fig. 7D). However, *pkr* transcript levels increased in Xela DS2 (Fig. 7E) and Xela VS2 (Fig. 7F) cells following poly(I:C) treatment. Xela DS2 showed a significant 5.8 ± 0.8 fold (p = 0.0079) increase in *pkr* transcript levels in response to poly(I:C) after 12 h, and this increase was similar at 24 h, with a 6.2 ±0.4 fold (p = 0.0079) increase above time matched non-treated controls (Fig. 7E). Transcripts for *pkr* were also up-regulated in a time-dependent manner in poly(I:C) treated Xela VS2 cells at all time points, reaching a 4.3 ± 0.7 fold increase (p = 0.0079) in *pkr* mRNA levels after 24 h post treatment (Fig. 7F). These data suggest that Xela DS2 and Xela VS2 may be secreting type I interferons in response to extracellular poly(I:C) that leads to the downstream upregulation of select ISGs and potential activation of antiviral programs.

### 3.6. Treatment of Xela DS2 and Xela VS2 cells with poly(I:C) induces the up-regulation of key proinflammatory cytokines

To determine whether extracellular poly(I:C) recognition by frog skin epithelial cells results in promoting a proinflammatory response, transcript levels of key proinflammatory cytokines were assessed. Xela DS2 and Xela VS2 showed an increase in *tnf, il1b,* and *il8* mRNA levels following poly(I:C) treatment (Fig. 8). Xela DS2 demonstrated significant increases in *tnf* (Fig. 8A) and *il1b* (Fig. 8C) transcript levels of 78.0 ± 26.4 fold (p = 0.0079) and 22.3 ± 6.5 fold (p = 0.0079), respectively, at 3 h post poly(I:C) treatment, and these increases in transcript levels persisted over the 24 h time period examined. Xela VS2 cells displayed a 75.7 ± 21.7 fold (p = 0.0079) increase in *tnf* mRNA levels (p = 0.0079; Fig. 8B) and a 14.0 ± 0.1 fold (p = 0.0079) increase in *il1b* mRNA levels (p = 0.0079; Fig. 8D) 6 h post poly(I:C) treatment. Transcript levels of *tnf* and *ilb* remained significantly elevated in Xela VS2 relative to the time-matched, non-treated controls over the entire course of the experiment, with a 146.7 ± 64.7 fold (p = 0.0079) increase in *tnf* expression (Fig. 8B) and a 26.2 ± 5.8 fold (p = 0.0079) increase in *il1b* expression (Fig. 8D) observed at 24 h post poly(I:C) treatment.

**Figure 8.**
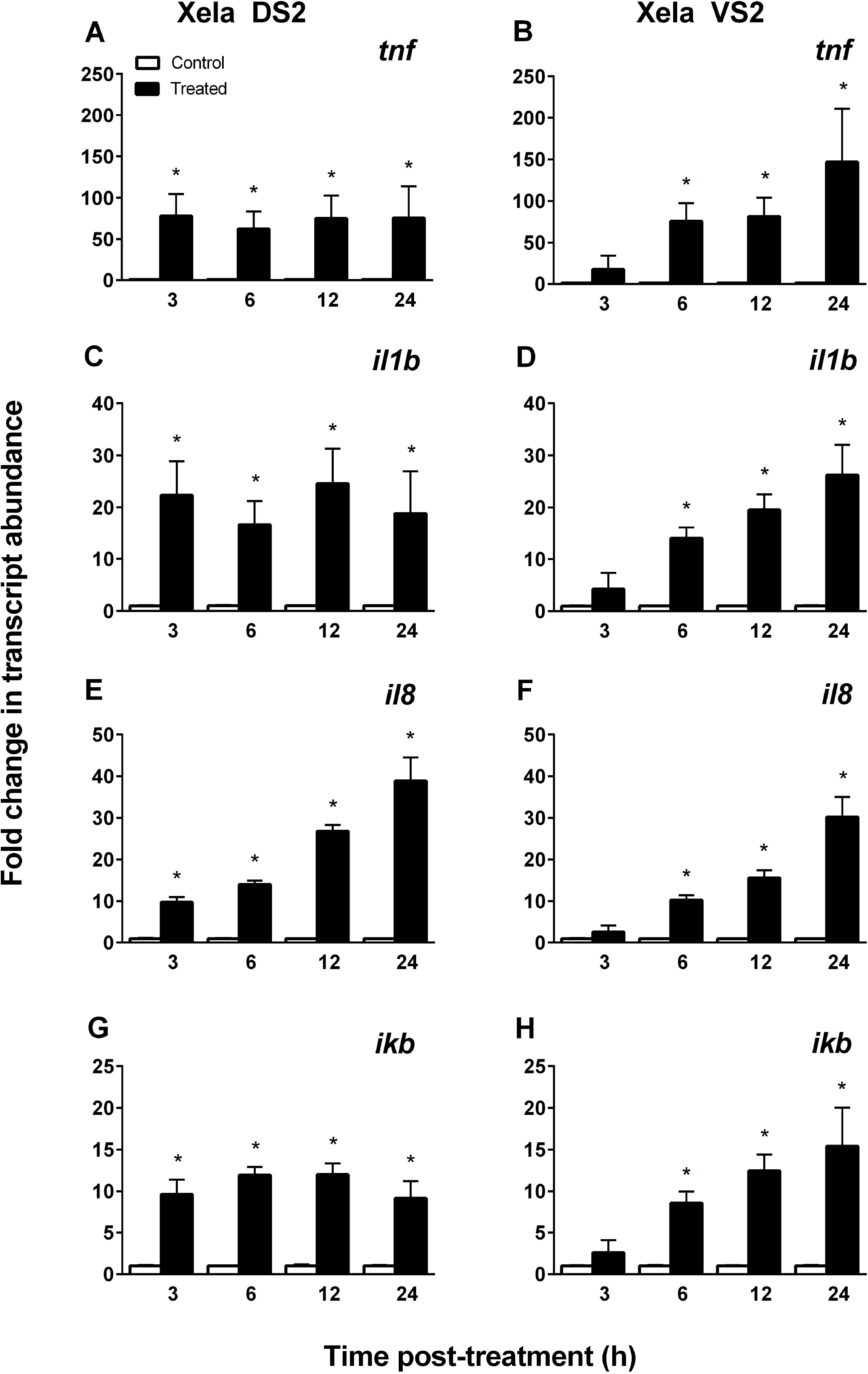
Poly(I:C) treatment induces the upregulation of cytokine and chemokine transcript levels in Xela DS2 and Xela VS2. RT-qPCR was performed on cDNA generated from non-treated (control) and poly I:C treated Xela DS2 or Xela VS2 cells to determine relative transcript levels of *tnf* (A, B), *il1b* (C, D), *il8* (E, F), *and ikb* (G, H). For all experiments, mRNA levels from treated samples were normalized to that of the respective time-matched controls, (n = 5 independent trials). Significant differences were determined by a Mann-Whitney test with Bonferroni correction (*p* < 0.0125).

As IL-8 is a potent chemokine and thus instrumental in recruiting innate immune cells (e.g. neutrophils) to the site of inflammation, we examined the relative change in *il8* transcripts in Xela DS2 and Xela VS2 cells in response to poly(I:C) stimulation. Similar to the upregulation of proinflammatory cytokine gene expressions, a time-dependent increase in *il8* transcript levels were observed in Xela DS2 (Fig. 8E) and Xela VS2 (Fig. 8F) as early as 3 h and 6 h post poly(I:C) treatment, respectively. By 24 h post treatment, *il8* transcript levels reached 38.9 ± 5.6 fold (p = 0.0079) and 30.1 ± 4.9 fold (p = 0.0079) higher levels in poly(I:C) treated Xela DS2 (Fig. 8E) and Xela VS2 (Fig. 2.8F), respectively, relative to time-matched, non-treated control cells.

Finally, changes in *ikb* expressions in Xela DS2 and Xela VS2 following treatment with poly(I:C) were examined. Xela DS2 showed a significant 9.6 ± 1.8 fold (p = 0.0079) increase in *ikb* transcript levels by 3 h post treatment and these transcript levels were sustained throughout the 24 h treatment period (Fig. 8G). Similarly, a significant 8.6 ± 1.4 fold (p = 0.0079) increase in *ikb* transcript levels was observed in Xela VS2 cells by 6 h post treatment, reaching 15.4 ± 4.6 fold higher transcript levels (p = 0.0079), relative to the time-matched controls, by 24 h post treatment (Fig. 8H).

## 4. Discussion

We hypothesized that frog skin epithelial cells would initiate proinflammatory and antiviral programs in response to poly(I:C), a synthetic analogue of viral dsRNA, and thus act as important early cellular sensors in initiating and directing frog skin innate immune responses. The goals of this study were to first generate frog skin epithelial cell lines from *X. laevis* and then to understand innate immune gene regulation in response to poly(I:C) treatment. This is the first reported study, to our knowledge, that has generated frog skin epithelial cell lines from *X. laevis* and examined their proinflammatory and antiviral gene expression responses to poly(I:C). The successful generation and use of two *X. laevis* skin epithelial-like cell lines from dorsal and ventral skin further allowed us to compare cellular responses in skin epithelial-like cells *in vitro* that may be indicative of the physiological function(s) of skin tissues *in vivo*.

### 4.1. Xela DS2 and Xela VS2 are skin epithelial-like cell lines

Xela DS2 and Xela VS2 have been developed from the dorsal and ventral skin of adult African clawed frogs and are the first known skin epithelial-like cell lines to be produced from the adult of this important model organism. Although a number of cell lines have been developed from *X. laevis*, they are mainly derived from embryos (Sakaguchi et al., 1989; Smith et al., 2002) or tadpoles (Pudney et al., 1973), with only a handful of cell lines developed from adult frogs, mostly derived from tumors (Rafferty, 1969; Robert et al., 1994) or restricted to kidney tissue origin such as the A6 (Rafferty, 1969) and XLK-WG (Martin et al., 1998) cell lines. Like other ectothermic vertebrate cell lines (Bols et al., 2017; Rafferty, 1969; Rausch and Simpson, 1988), Xela DS2 and Xela VS2 arose spontaneously through subculturing efforts and have been passaged over 160 times within the past four years. Both Xela DS2 and Xela VS2 grow optimally at 26°C; however, they are capable of growth over a much wider temperature range (14°C to 30°C) and thus excellent models for temperature-based research. Xela DS2 and Xela VS2 have an absolute requirement for FBS, generally exhibiting increased growth with increasing FBS concentration. Although Xela DS2 and Xela VS2 initially exhibited low levels of senescence associated beta-galactosidase activity at early passages, beta-galactosidase activity was completely absent at higher passages. In mammalian cells, detection of beta-galactosidase activity at a pH of 6 is usually associated with cellular senescence and is rarely observed in quiescent or immortalized cells (Dimri et al., 1995), permitting beta-galactosidase activity to be used as a general indicator of cellular senescence *in vivo* and *in vitro* (Itahana et al., 2007). The long-term and high proliferative capacity, together with the lack of beta-galactosidase activity, supports that Xela DS2 and Xela VS2 demonstrate inherent characteristics of immortalized cells.

Xela DS2 and Xela VS2 display uniform epithelial-like cell morphology of frog epidermal cells in culture (Nishikawa et al., 1990) and express epithelial cell associated molecular markers characteristic of epithelial cells of humans and mice. Both Xela DS2 and Xela VS2 express the *cdh1* gene wherein the corresponding cadherin 1 protein, also known as epidermal cadherin (E-cadherin), is involved in the formation of adherens junctions specifically between epithelial cells (Proksch et al., 2008; van Roy and Berx, 2008). Since E-cadherin is a known molecular marker in epithelial tissues for cell-to-cell adhesion, the expression of *cdh1* in Xela DS2 and Xela VS2 suggest that these cells are epithelial-like. These cells also expressed *cldn1* and *ocln* genes necessary for the formation of tight junctions in a number of cell types, including epithelial cells (Tsukita and Furuse, 1999). The presence of claudin-1 is important in maintaining mammalian epidermal barrier integrity and barrier function, evident by the death of claudin-1-deficient mice, 1 d after birth (Furuse et al., 2002). Further evidence that Xela DS2 and Xela VS2 may be epithelial in nature is the expression of cytokeratin gene markers. Although Xela DS2 and Xela VS2 demonstrate differential *krt5* expression and complete lack of *krt14* expression, keratins which are expressed in complex epithelium, both cell lines express *krt19*, a keratin subtype known to be expressed in highly permeable simple epithelium (Chu and Weiss, 2002). This coincides with the characteristic permeable nature of frog skin, as the epidermal layer is relatively thin compared to skin of other vertebrates in order to achieve important physiological functions (Lillywhite, 2006). Furthermore, cytokeratin 19 has been implicated as a marker for skin stem cells that are necessary for continuous regeneration of skin layers and wound healing, and the presence of cytokeratin 19 has been identified in human skin cultures *in vitro* (Michel et al., 1996). However, it is important to acknowledge that expressions of cytokertains and vimentins in epithelial cells from different tissues and species (e.g. fish) vary widely (e.g. Markl and Franke, 1988) and thus it is challenging to confirm the epithelial nature of Xela DS2 and Xela VS2 in the absence of well characterized expression of these markers in frog skin epithelial cells *in vivo*.

Interestingly, Xela DS2 and Xela VS2 exhibit differential expression of *col* and *vim* molecular markers. Collagen genes are usually expressed by fibroblasts and the production of collagen proteins is important in the formation of the extracellular matrix (Groulx et al., 2011). However, collagen genes are also expressed in epithelial cells (Creely et al., 1988), including mucosal-associated epithelial cells (Birkedal-Hansen et al., 1982) or epithelial cells undergoing EMT (Hosper et al., 2013). It is possible that a select population of Xela DS2 and Xela VS2 cells may be undergoing EMT as *vim*, a gene commonly expressed by mesenchymal cells (Chaw et al., 2012), was expressed in both cell lines. While our results indicate that Xela DS2 and Xela VS2 differentially express collagen genes, the synthesis and secretion of collagen proteins and formation of extracellular matrices by these cells lines requires further investigation. The expression of tight and adherens junction transcripts, particularly *cdh1*, is strong evidence to support that Xela DS2 and Xela VS2 are epithelial-like in nature, as cellular junctions are important for maintaining epidermal integrity and *cdh1* is known to be localized to epidermal cells in epithelial tissues (Farquhar and Palade, 1963; Proksch et al., 2008; van Roy and Berx, 2008). Collectively, these results, in conjunction with cell morphology, strongly support that Xela DS2 and Xela VS2 are epithelial-like in origin, although further *in vivo* detection of these epithelial markers at the protein level is needed to confirm their presence. While the expression of junction markers, filaments (cytokeratins and vimentins), and collagen in these cells appear to align with observations in mammalian systems, it is important to acknowledge the inherent differences between amphibian and mammalian models due to the complexity of each respective system in tandem with the lack of studies characterizing the expression of these genes/proteins in adult amphibians. Nonetheless, the establishment and characterization of Xela DS2 and Xela VS2 represent a key advancement in the generation of *in vitro* model systems for future investigations in amphibian skin epithelial cell biology, including environmental parameters that may impact epithelial cell characteristics such as cellular junctions and transepithelial resistance as a means of determining skin barrier integrity.

### 4.2. Xela DS2 and Xela VS2 initiate proinflammatory and antiviral programs in response to extracellular poly(I:C)

Xela DS2 and Xela VS2 appear highly sensitive to extracellular poly(I:C) based on early (within 3 h) changes in cell morphology that resulted in a proportion of cells contracting, rounding, and losing adherence to the culture vessel. While microgram per mL quantities of poly(I:C) are widely used to treat animal cells without reported changes to cell morphology or viability (Holopainen et al., 2012; Kumar et al., 2006), cell cytotoxicity in response to poly(I:C) has been observed in some cultured animal cells, indicating response to poly(I:C) is dose, cell type and species dependent. Across the limited studies examining response of amphibian cultured cells to poly(I:C) treatment, the *X. laevis* A6 kidney epithelial cell line was not reported to exhibit changes in cell morphology or viability (Sang et al., 2016), while poly(I:C)-induced cytotoxicity was observed in a tadpole cell line from American toad (*Anaxyrus americanus*, previously *Bufo americanus*) (Vo et al., 2019). Similarly, poly(I:C) is not reported to affect cell morphology or viability in some cultured fish cells (Abram et al., 2019; Semple et al., 2018) but is cytotoxic to others, such as walleye (Abram et al., 2020). Despite the variability in cellular response to extracellular poly(I:C) treatment, the mechanisms underlying the variance in poly(I:C) induced cytotoxicity across cell types and species are unclear. It is interesting to consider whether the poly(I:C)-induced loss of cellular adherence in Xela DS2 and Xela VS2 may be linked to the increased skin sloughing observed in some frog species in response to pathogens (Ohmer et al., 2017). However, given the limited studies on amphibian skin tissue and epithelial cell responses to extracellular viral dsRNA, pathogens in general, and the mechanisms that drive skin sloughing in response to pathogens, further investigation is required to ascertain whether our *in vitro* observations in Xela DS2 and Xela VS2 following poly(I:C) treatment are reflective of in *vivo* skin sloughing responses.

Concurrent with changes in cell morphology, poly(I:C)-treated Xela DS2 and Xela VS2 demonstrated marked increases in mRNA levels for key antiviral (*ifn1*) and proinflammatory (*il1b*, *tnf*) genes, likely through IRF and NF-κB dependent gene regulatory pathways, that are known to encode for proteins essential in activating antiviral and proinflammatory responses. In addition, the time-dependent increase in *il8* mRNA levels in poly(I:C)-treated Xela DS2 and Xela VS2 suggest that frog skin epithelial cells may play an important role in recruitment of immune cells to the skin during viral infections. Induction of antiviral responses were assessed by examining the expression of an intron-containing type I IFN that most closely resembles *ifn6* in the expanded *X. laevis* type I IFN gene family repertoire (Sang et al., 2016). The type I IFN group, which includes the *ifn6* gene, was found to be poly(I:C)-responsive, suggesting it is important in mediating antiviral responses (Sang et al., 2016). Similarly, recombinant *X. laevis* type I IFN was shown to induce ISGs to mediate antiviral programs (Grayfer et al., 2015; Wendel et al., 2017). While we did not have access to an antibody against *X. laevis* IFN to assess secreted IFN, the increases in *pkr* transcript levels that followed the modest increase in *ifn1* transcripts in both cell lines suggest that type I interferons were likely being secreted and binding to their cognate receptors to upregulate ISGs known to play an integral role in the inhibition of viral replication in mammalian models (Schoggins and Rice, 2011). Interestingly, we observed a more robust increase in transcripts for proinflammatory genes than the type I interferon gene we examined in poly(I:C) treated Xela DS2 and Xela VS2. However, the biological significance of this observation requires further investigation. Collectively, our findings provide evidence that Xela DS2 and Xela VS2 respond to extracellular poly(I:C) to initiate antiviral programs. Investigations of functional antiviral responses of Xela DS2 and Xela VS2 to viruses affecting frog health are currently underway in our laboratory.

To our knowledge, this study is the first to demonstrate adult frog skin epithelial cells are capable of responding to extracellular viral dsRNA, albeit a synthetic analogue. Studies in human, fish, and more recently in frog/tadpole models have shown the importance of SR-As in extracellular dsRNA recognition and subsequent trafficking to endosomes where it can be recognized by TLR3 to initiate downstream intracellular signaling pathways (DeWitte-Orr et al., 2010; Matsumoto and Seya, 2008; Poynter and DeWitte-Orr, 2018; Vo et al., 2019). Although we did not examine the presence of SR-As, the downstream antiviral and proinflammatory gene regulation in response to extracellular poly(I:C) in Xela DS2 and Xela VS2 suggest that these cells possess SR-As that are necessary for viral dsRNA trafficking to TLR3 in endosomes. Further investigation is required to determine the profile of SR-As and TLRs expressed in Xela DS2 and Xela VS2.

## 5. Conclusion

In conclusion, we have generated novel skin epithelial-like cell lines derived from adult *X. laevis* dorsal and ventral skin tissues. Xela DS2 and Xela VS2 exhibit differences in growth and expression of molecular markers and are capable of responding to extracellular poly(I:C) resulting in potent upregulation of antiviral and proinflammatory genes, albeit with differing kinetics. Our findings provide evidence of the conservation of the ability of epithelial cells across vertebrates to recognize and respond to dsRNA. The ability of Xela DS2 and Xela VS2 to initiate antiviral and proinflammatory mechanisms suggest that frog skin epithelial cells may be more than just an inert physical barrier and instead be active participants in initiating and directing innate immune responses to viral infections in the skin. This implies that frog skin epithelial cells are thus an important first line of defence in sensing pathogens and signalling to underlying immune cells that a threat is present. Furthermore, the ability for frog skin epithelial cells to respond to viral dsRNA may be important for promoting frog immunocompetency by limiting the spread of viral infection. Overall, Xela DS2 and Xela VS2 may serve as powerful *in vitro* systems for studying amphibian host-pathogen interactions at the skin epithelial interface in tandem with the impact of our changing environment and may prove novel and valuable tools to the field of amphibian skin biology.

## 7. Acknowledgements

The authors thank Dr. Mungo Marsden for the kind donation of *Xenopus laevis* tissues from which these cell lines were derived. This work was supported by a Natural Sciences and Engineering Research Council (NSERC) Discovery Grant (RGPIN-2017-04218) to BAK. MPB and JFAV received financial support in the form of Graduate Teaching Assistantships, Science Graduate Experience Awards and Science Graduate Student Awards from the Department of Biology and University of Waterloo.

## Figure Legends

**Supplemental Figure 1. Multiple sequence alignment of Xela DS2 COI clone sequences against reference COI region of *X. laevis* mitochondrial genome.** Clustal Omega sequence alignment of sequenced COI clones expressed in Xela DS2 against reference *X. laevis* mitochondrial genome (GenBank Accession HM991335.1) at the COI region (nucleotide position 5396-6952, indicated by ►), viewed through Jalview software. Shading pattern indicates percentage identity across sequences, wherein like shading within columns indicates sequence matching.

**Supplemental Figure 2. Multiple sequence alignment of Xela VS2 COI clone sequences against reference COI region of *X. laevis* mitochondrial genome.** Clustal Omega sequence alignment of sequenced COI clones expressed in Xela VS2 against reference *X. laevis* mitochondrial genome (GenBank Accession HM991335.1) at the COI region (nucleotide position 5396-6952, indicated by ►), viewed through Jalview software. Shading pattern indicates percentage identity across sequences, wherein like shading within columns indicates sequence matching.

**Supplemental Figure 3. Multiple sequence alignment of Type I IFN sequences.** Clustal Omega sequence alignment of sequenced type I IFN expressed in Xela DS2 and Xela VS2 (indicated by ►) against reference type I IFN (GenBank Accession KF597522.1) where original RT-qPCR primers were designed from (indicated by ●), and the expanded type I IFN repertoire including *ifn1* (KU594561.1), *ifn2* (KU594562.1), *ifn3* (KU594563.1), *ifn4* (KU594564.1), *ifn5* (KU594565.1), *ifn6* (KU594566.1), and *ifn7* (KU594567.1), viewed through Jalview software. Shading pattern indicates percentage identity across sequences; like shading within columns indicates sequence matching and differential shading across columns indicates sequence variability wherein darker shades have the least variability.

