## Supplementary figures and images for "Xela DS2 and Xela VS2: two novel skin epithelial-like cell lines from adult African clawed frog (*Xenopus laevis*) and their response to an extracellular viral dsRNA analogue"

### Suppl. Fig. 1

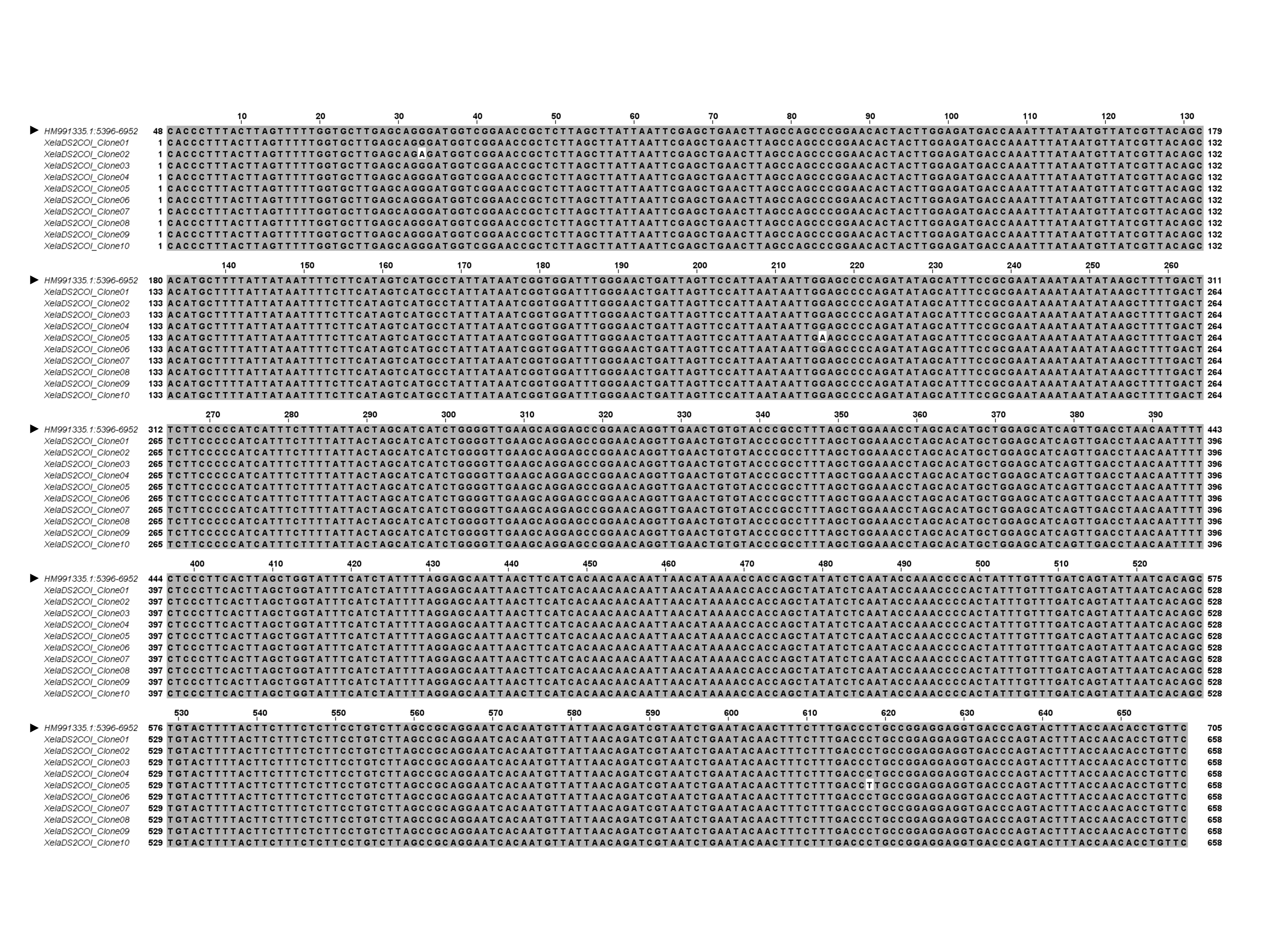

### Suppl. Fig. 2

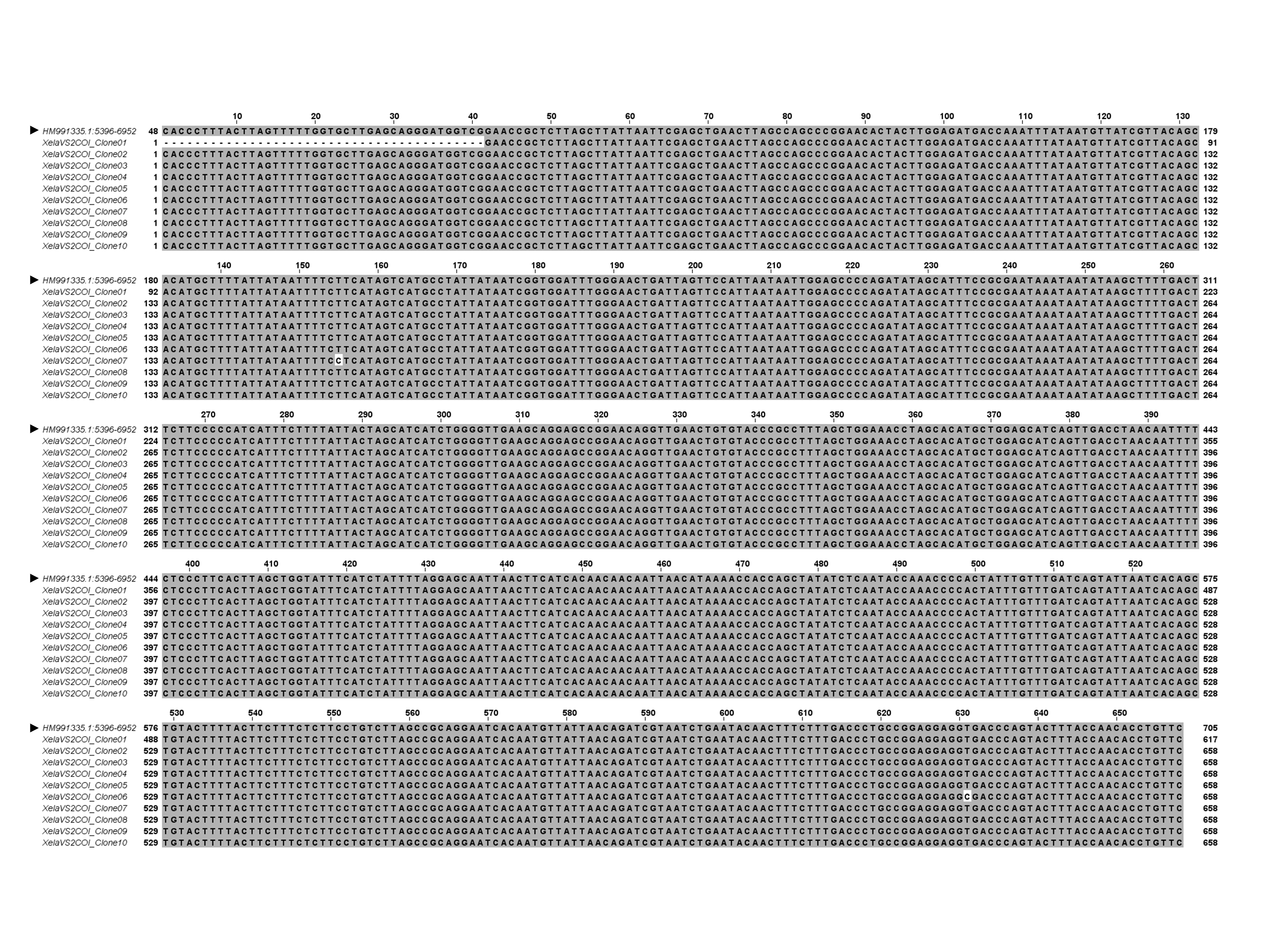

### Suppl. Fig. 3

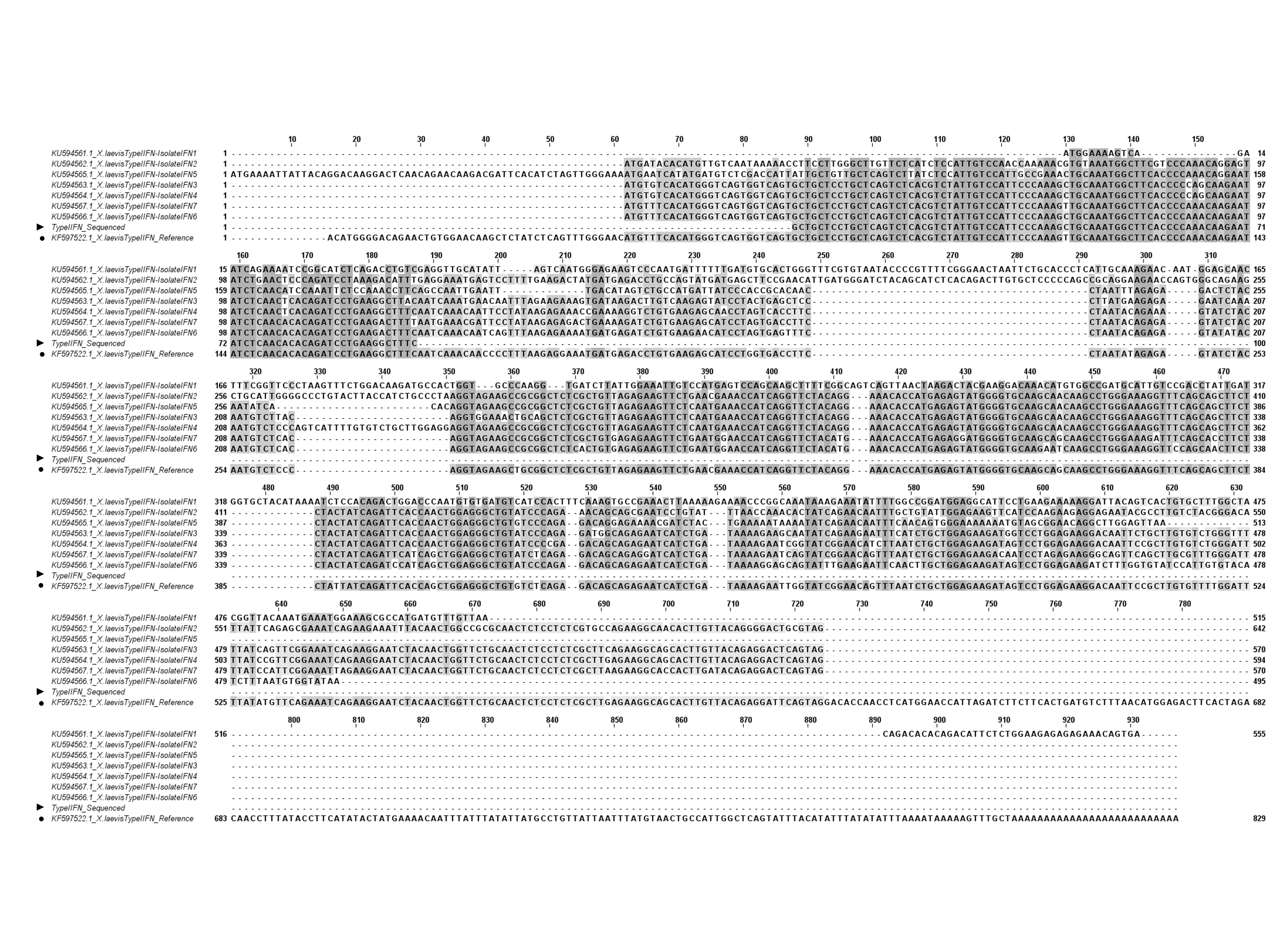
