## Supplementary material for "Xela DS2 and Xela VS2: two novel skin epithelial-like cell lines from adult African clawed frog (*Xenopus laevis*) and their response to an extracellular viral dsRNA analogue": Suppl. Table 1

**Supplemental Table 1. Candidate endogenous genes for use in RT-qPCR analysis of poly(I:C) treated Xela DS2 and Xela VS2 cells.**

| **Target** | **Xela DS2  M-Value** | **Xela VS2  M-Value** |
| --- | --- | --- |
| *actb* | 0.528 | 0.266 |
| *cyp* | 0.354 | 0.226 |
| ***ef1a*** | **0.278** | **0.217** |
| *gapdh* | 0.290 | 0.348 |
| *hgprt* | 0.367 | 0.298 |
